# Culture conditions greatly impact the levels of vesicular and extravesicular Ago2 and RNA in extracellular vesicle preparations

**DOI:** 10.1101/2023.06.21.545797

**Authors:** Lizandra Jimenez, Bahnisikha Barman, Youn Jae Jung, Lauren Cocozza, Evan Krystofiak, Cherie Saffold, Kasey C. Vickers, John T. Wilson, T. Renee Dawson, Alissa M. Weaver

**Affiliations:** Department of Cell and Developmental Biology, Vanderbilt University School of Medicine, Nashville, Tennessee; Center for Extracellular Vesicle Research, Vanderbilt University School of Medicine, Nashville, Tennessee; Department of Chemical and Biomolecular Engineering, Vanderbilt University School of Engineering, Nashville, Tennessee; Cell Imaging Shared Resource EM Facility, Vanderbilt University, Nashville, Tennessee; Department of Pathology, Microbiology and Immunology, Vanderbilt University Medical Center, Nashville, Tennessee; Department of Medicine, Vanderbilt University Medical Center, Nashville, Tennessee

**Keywords:** extracellular RNA, extracellular vesicles, Argonaute 2, serum, RNA-binding proteins

## Abstract

Extracellular vesicle (EV)-carried miRNAs can influence gene expression and functional phenotypes in recipient cells. Argonaute 2 (Ago2) is a key miRNA-binding protein that has been identified in EVs and could influence RNA silencing. However, Ago2 is in a non-vesicular form in serum and can be an EV contaminant. In addition, RNA-binding proteins (RBPs), including Ago2, and RNAs are often minor EV components whose sorting into EVs may be regulated by cell signaling state. To determine the conditions that influence detection of RBPs and RNAs in EVs, we evaluated the effect of growth factors, oncogene signaling, serum, and cell density on the vesicular and nonvesicular content of Ago2, other RBPs, and RNA in small EV (SEV) preparations. Media components affected both the intravesicular and extravesicular levels of RBPs and miRNAs in EVs, with serum contributing strongly to extravesicular miRNA contamination. Furthermore, isolation of EVs from hollow fiber bioreactors revealed complex preparations, with multiple EV-containing peaks and a large amount of extravesicular Ago2/RBPs. Finally, KRAS mutation impacts the detection of intra- and extra-vesicular Ago2. These data indicate that multiple cell culture conditions and cell states impact the presence of RBPs in EV preparations, some of which can be attributed to serum contamination.

## Introduction

Extracellular vesicles (EVs) are small lipid-bound particles that carry a diverse array of cargos and facilitate intercellular communication (Dixson et al., 2023; Maas et al., 2017). EVs are released from cells by many mechanisms and include small EVs (≤150 nm) and large EVs (≥150 nm). Small EVs can be formed as intraluminal vesicles (ILVs) within endosomal multivesicular bodies (MVBs) that then are secreted as exosomes from the cells upon fusion with the plasma membrane. Small EVs can also bud from the plasma membrane (Dixson et al., 2023; Maas et al., 2017). EV cargoes include proteins, lipids, metabolites, and nucleic acid species and are thought to play an important role in driving cancer progression and aggressiveness (Dixson et al., 2023; Maas et al., 2017; Sato & Weaver, 2018). EVs have been shown to affect multiple cellular processes, including cancer cell migration, invasion, metastasis, proliferation, angiogenesis, and drug resistance (Aucher et al., 2013; Chen et al., 2014; Dhondt et al., 2016; Jiang et al., 2023; Liao et al., 2016; Sahoo et al., 2011; Zhang et al., 2010).

RNA EV cargoes, such as miRNAs, have the potential to control gene expression of recipient cells and are of both biologic and therapeutic interest (Dixson et al., 2023; O’Brien et al., 2020). Some of the functions of RNAs transferred to recipient cells include promotion of cell proliferation, tumor growth, angiogenesis, cancer metastasis, drug resistance, and modulation of immune activation (Chiou et al., 2018; Dixson et al., 2023; Li et al., 2021; Patton et al., 2015). EV-mediated delivery of therapeutic RNAs such as siRNAs or mRNAs is being actively explored for treatment of human disease (Cecchin et al., 2023; Murphy et al., 2019).

Despite the potential importance of EV-carried RNAs, there is limited knowledge regarding how sorting of RNA into EVs occurs. Numerous studies have shown that RNA trafficking into EVs depends on a variety of RNA-binding proteins (RBPs), including hnRNP A2/B1, SYNCRIP, YBX-1, and Argonaute-2 (Ago2) (Hobor et al., 2018; Liu et al., 2021; McKenzie et al., 2016; Santangelo et al., 2016; Villarroya-Beltri et al., 2013; Zhou et al., 2020). Sorting of these RNA-RBP complexes may occur at ER-membrane contact sites and depend on ceramide synthesis and the autophagy protein LC3 (Barman et al., 2022; Dixson et al., 2023; Leidal et al., 2020; O’Brien et al., 2020).

Ago2 is a key component of the RNA-induced silencing complex (RISC) complex, having slicer activity for miRNA-associated mRNA (Meister, 2013). In addition, Ago2 binds to tRNAs, ribosomal RNAs, snoRNAs and vault RNAs (Atwood et al., 2016; Burroughs et al., 2011; Dupuis-Sandoval et al., 2015; Haussecker et al., 2010; Kishore et al., 2013; Persson et al., 2009; Woolnough et al., 2015); all of these are RNA species that are found in EVs (Lunavat et al., 2015; Shurtleff et al., 2016; van Balkom et al., 2015). Due to the potential for Ago2 to control gene downregulation in recipient cells and transport a variety of small RNAs, we and others have studied its presence and trafficking into EVs (McKenzie et al., 2016; Weaver & Patton, 2020). We previously found that Ago2 is present on the inside of EVs and that its trafficking into small EVs is regulated by KRAS-MEK-ERK signaling (McKenzie et al., 2016) and ER-membrane contact site formation (Barman et al., 2022). However, other groups have found that Ago2 is primarily an extravesicular contaminant of EVs (Arroyo et al., 2011; Jeppesen et al., 2019; Turchinovich et al., 2011) and that RNA is a minor component of EVs (Albanese et al., 2021). These observed differences between studies can be partially explained by the effect of oncogenic mutations driving signaling (e.g., KRAS) in the cell line from which EVs were isolated. Another major point of divergence among many studies is the differential use of serum in conditioning media, which contains contaminating EVs and a large amount of nonvesicular Ago2 and RNA (Arroyo et al., 2011; Driedonks et al., 2019; Kornilov et al., 2018; Mannerstrom et al., 2019; Shelke et al., 2014; Thery et al., 2006; Turchinovich et al., 2011; Wei et al., 2016). Current MISEV guidelines recommend ultracentrifugation to deplete EVs from FBS prior to use, yet this process is only 70-80% effective and leaves substantial numbers of residual EVs and lipoproteins in the “EV-depleted” serum (Lehrich et al., 2018; Pham et al., 2021).

Beyond media contaminants, growing evidence suggests that many aspects of cell culture environments can introduce profound differences in the RNA and protein cargoes of isolated EVs (Casajuana Ester & Day, 2023; Urzi et al., 2022). The use of 3D culturing approaches has revealed significant differences in the levels of miRNAs and other small RNAs found in EVs compared to those isolated from monolayer cell culture (Han et al., 2022; Thippabhotla et al., 2019; Tu et al., 2021; Yuan et al., 2022). In addition to the extracellular RNA and RBP contaminants found in fetal bovine serum (FBS), the presence of FBS in conditioning media introduces a mixture of growth factors, hormones, cytokines and nucleic acids that differ between serum lots and could alter EV biogenesis in undefinable ways (Zheng et al., 2006). The use of defined media supplements in place of serum can help to minimize these concerns; however, many cell lines do not tolerate this approach and defined supplements have also been shown to introduce contaminating RNA into EV preparations (Auber et al., 2019). Overall, there is ample reason to carefully weigh the potential for extraneous influences of media, culturing conditions, and genetic variation among cell lines on EVs isolated from cell culture and the evaluation of their RNA and protein cargoes.

In this study, we set out to comprehensively evaluate the effects of diverse cell culture conditions on SEV character and RNA/RBP content, carefully resolving the association of Ago2, other RBPs and miRNAs with specific vesicular and non-vesicular populations across high resolution density gradients. We compared multiple cell culture and cell state conditions, including comparing EVs from colon cancer cells containing wildtype or mutant KRAS alleles, comparison of three different culture media conditions (serum-free DMEM, DMEM supplemented with EV-depleted fetal bovine serum (FBS), and Opti-MEM), and 2D flask culture vs. high density culture in hollow fiber bioreactors. Under 2D flask culture conditions, we found that serum is indeed a prominent source of nonvesicular Ago2 and RNA. In addition, conditioning the cells in serum-free DMEM led to higher levels of vesicular RBPs than DMEM supplemented with EV-depleted FBS or Opti-MEM, which contains growth factors but no serum. We also found that highly purified preparations of EVs derived from serum-containing conditioned media collected from 2D flask culture were contaminated with a significant amount of extravesicular RNA. Finally, density gradient fractionation of either serum-free or serum-containing conditioned media collected from hollow fiber bioreactors revealed multiple EV peaks and substantial amounts of nonvesicular Ago2 and other RBPs that were not derived from the media. These data indicate that cell signaling and context regulate the RBP and RNA content of EVs. Furthermore, contaminating nonvesicular RNA-RBP complexes from serum or released from cells may greatly complicate interpretation of results and necessitate rigorous purification and analysis procedures.

## Materials and methods

### Cell culture

DKs-8 and DLD-1 cells (Cha et al., 2015; Shirasawa et al., 1993) were cultured in DMEM (cat no. 10-013-CV, Corning) supplemented in 10% fetal bovine serum (FBS, cat no. F0926, Sigma), non-essential amino acids (NEAA, cat no. 25-025-CI, Corning), and L-Glutamine (cat no. 25030081, Gibco). EV-depleted FBS was made by ultra-centrifuging the FBS at 100,000 x g overnight (18h) in a Ti 45 rotor (Beckman Coulter).

### Isolation of EVs from conditioned medium

For flask cultures, DKs-8 or DLD-1 cells were plated in DMEM growth media. The next day, the cells were washed three times with 1x PBS (cat no. 21-040-CV, Corning). Serum-free DMEM (DMEM supplemented with L-glutamine and NEAA), 10% EV-depleted FBS in DMEM, Opti-MEM, or Serum-free DMEM with 100 ng/mL epithelial growth factor (EGF) (Larios et al., 2020; Oszvald et al., 2020) were added to the cells. After 48 hours, the conditioned medium was collected from the cells and the EVs were isolated via serial centrifugation. Floating live cells and dead cell debris were removed from the conditioned medium after centrifugation steps of 300 x g for 10 minutes and 2,000 x g for 25 minutes, respectively. Large EVs were collected by centrifugation at 10,000 x g for 30 minutes in a Ti 45 rotor (Beckman Coulter). Small EVs were collected by a modified cushion-density gradient protocol (Li et al., 2018; Sato et al., 2019). The following modifications were made: 1) the supernatant from 10,000 x g spin was concentrated with Centricon Plus-70 centrifugal filter (cat no. UFC710008, Millipore), per manufacturer’s instructions. 2) The concentrate was overlaid onto 2 mL of 60% iodixanol (OptiPrep) and centrifuged at 100,000 x g either for 4 hr (hollow fiber bioreactor) or overnight (2D flask culture) in a SW32 rotor (Beckman Coulter). For the preparation of the density gradient, the 40% iodixanol layer was made by transferring the bottom 3 mL of the cushion (which includes 1 mL of collected EVs and 2 mL of 60% iodixanol) into a new tube. 3 mL of 20% iodixanol, 10% iodixanol, and 5% iodixanol were subsequently layered on top. These iodixanol dilutions were prepared by diluting the OptiPrep with 0.25 M sucrose/10 mM Tris, pH 7.5. After an 18-h centrifugation step at 100,000 x g in a SW40 rotor (Beckman Coulter), 12 density gradient fractions were collected, diluted in PBS, and centrifuged at 100,000 x g for 3 h. In Figure 1, small EVs were also collected by centrifugation at 120,000 x g for 4 hr, which was subsequently bottom loaded to a density gradient (Jeppesen et al., 2019). To quantitate the size and concentration of the DKs-8 and DLD-1 LEVs and SEV-containing fractions, nanoparticle tracking analysis was performed using a Particle Metrix ZetaView PMX 110.

**Figure 1.**
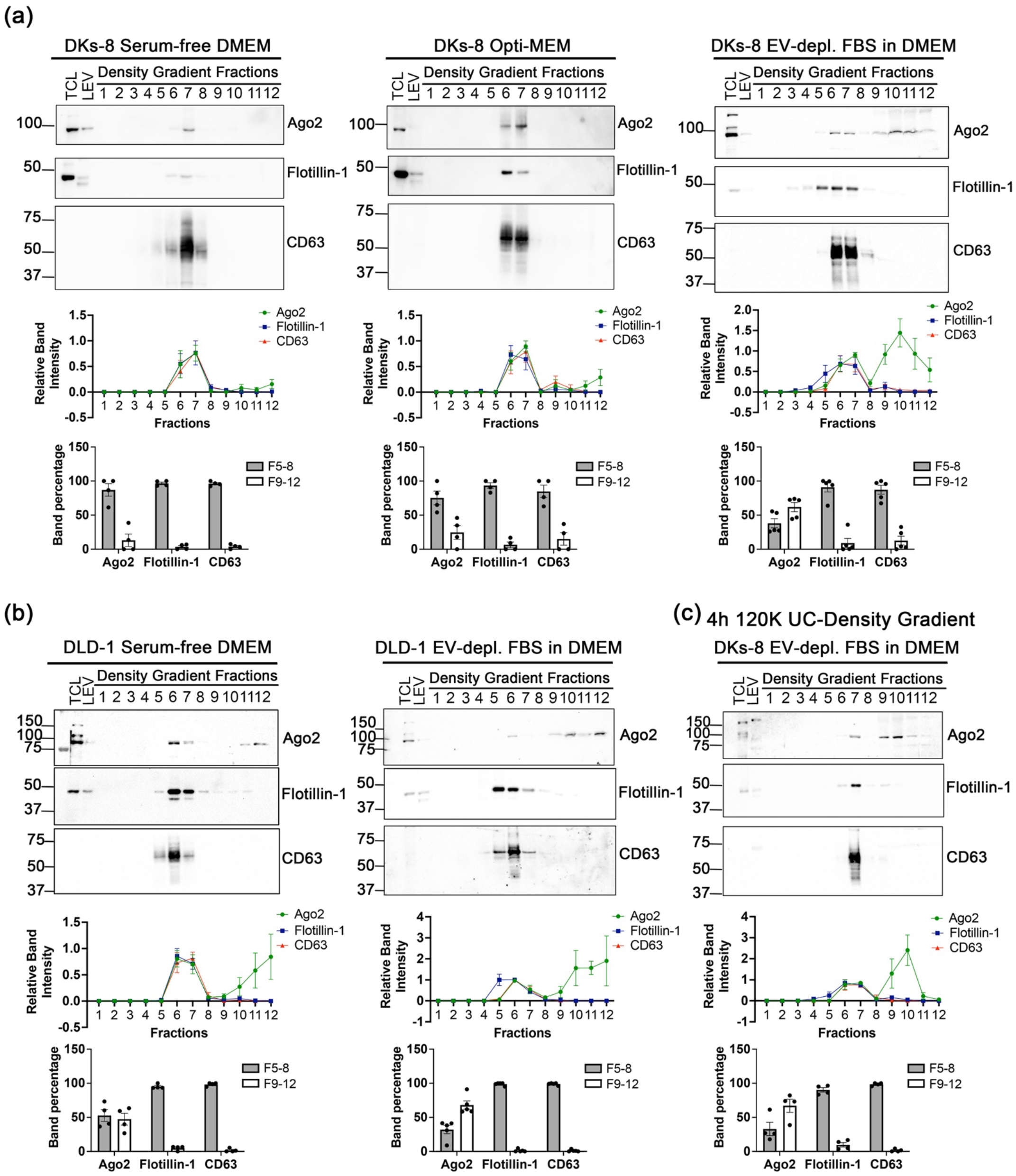
Ago2 is present as both vesicular and nonvesicular in EV-depleted serum. (a) Western blot analyses of DKs-8 TCL, LEV, and DG fractions from iodixanol cushions from the different media conditions probed for Ago2, Flotillin-1, and CD63. <u>Middle panel:</u> Relative intensity of the Ago2, Flotillin-1, and CD63 bands in the Cushion DG fractions from the different media conditions, normalized to the band intensity of the highest SEV lane (either Fraction 6 or 7) (n=4 or 5). <u>Bottom panel:</u> Percentage of protein bands found associated with Fractions 5-8 and 9-12. Data plotted as Mean and S.E.M. (b) Western blot analyses of DLD-1 TCL, LEV, and DG fractions from iodixanol cushions from the different media conditions probed for Ago2, Flotillin-1, and CD63. <u>Middle panel:</u> Relative intensity of the Ago2, Flotillin-1, and CD63 bands in the Cushion DG fractions from the different media conditions, normalized to the band intensity of the highest SEV lane (either Fraction 6 or 7) (n=4). <u>Bottom panel:</u> Percentage of protein bands found associated with Fractions 5-8 and 9-12. Data plotted as Mean and S.E.M. (c) Western blot analyses of DKs-8 TCL, LEV, and DG fractions from 4h UC to DG from EV-depleted FBS in DMEM condition probed for Ago2, Flotillin-1, and CD63. <u>Middle panel:</u> Relative intensity of the Ago2, Flotillin-1, and CD63 bands in the DG fractions from 4h UC to DG from EV-depleted FBS in DMEM condition, normalized to the band intensity of the highest SEV lane (either Fraction 6 or 7) (n=4). <u>Bottom panel:</u> Percentage of protein bands found associated with Fractions 5-8 and 9-12. Data plotted as Mean and S.E.M.

### Hollow fiber bioreactor cell culture

Prior to cell inoculation, cartridges were primed with sterile PBS for 24 h, followed by serum-free DMEM for 24 h and 10% FBS in DMEM with for 24 h. 2.5 x 10^8^ cells of both DKs-8 and DLD-1 were inoculated in separate hollow fiber cartridges with 20-kDa molecular weight cut-off (cat. no.: C2011, FiberCell Systems). Cells were maintained more than a week in 10% FBS in DMEM, non-essential amino acids, and L-glutamine. 10 µL of media was collected from the media to verify total glucose content using a glucose monitor (GlucCell^®^). Glucose uptake rate was calculated by taking the starting glucose level and subtracting it from the glucose level 24 h later and multiplying the volume of media in the media bottle. When the glucose uptake rate reached 1 g/day, cells were adapted by gradually increasing the proportion of chemically defined media for high density cell culture (CDM-HD) (cat no. CDM-HD, FiberCell Systems) or EV-depleted FBS up to 10% in DMEM (cat no. 10-013-CM). Conditioned medium (25 mL) was collected for harvest every 24 h for DLD-1 cells and 48 h for DKs-8 cells. Cell debris was removed by 300 x g for 5 min and 2,000 x g for 25 min centrifugation, and supernatants were stored at 4 °C for further purification.

### Antibodies

Rabbit anti-Argonaute 2 (cat no. 2897) was purchased from Cell Signaling. Rat anti-Ago2 (cat no. SAB4200085) and goat anti-Apolipoprotein B (cat no. AB742) were purchased from Millipore Sigma. Rabbit anti-CD63 (cat no. ab134045), rabbit anti-hnRNPA2B1 (cat no. ab183654), rabbit anti-hnRNP Q (SYNCRIP) (cat no. ab189405), rabbit anti-Tsg101 (cat no. 30871), and rabbit anti-YB1 (cat no. ab12148) were purchased from Abcam. Mouse anti-Flotillin-1 (cat no. 610820) was purchased from BD BioSciences. Mouse anti-Hsp70 (cat no. sc-66048) was purchased from Santa Cruz. Anti-Rabbit IgG (H+L), HRP Conjugated (cat no. W4011) and Anti-Mouse IgG (H+L), HRP Conjugate (cat no. W4021) were purchased from Promega. Goat anti-Rat IgG. (H+L), HRP Conjugated (cat no. 31470) and Rabbit anti-Goat IgG (H+L), HRP Conjugated (cat no. 31402) were purchased from Thermo Fisher Scientific.

### Western blot analysis

For Western blots, DKs-8 and DLD-1 cells were lysed in RIPA buffer (50 mM Tris pH 7.6, 150mM NaCl, 1% SDS, 1% NP-40, and 0.5% Sodium Deoxycholate) with Protease Inhibitor cocktail (cat no. 046693124001, Roche) and 1μM PMSF (cat no. 44-865-0, Tocris Bioscience). The protein concentrations of total cell lysates (TCL) were determined utilizing Pierce BCA Assay (cat no. 23225, Thermo Fisher Scientific). The protein concentrations of the EVs were determined utilizing Pierce Micro BCA Assay (cat no. 23235, Thermo Fisher Scientific). For Western blots of density gradient fractions, 10 μg of DKs-8 or DLD-1 TCLs and LEVs and half of the resuspended gradient fractions or were boiled in 2x SDS-PAGE sample buffer (4x SDS sample buffer: 0.2M Tris, pH 6.8, 8% SDS, 40% glycerol, 0.16M DTT and 0.29M bromophenol blue in water) for 5 min and loaded on 15-well 9% polyacrylamide gels or NuPage 3-8% Tris - Acetate gels (ApoB-100 WBs). SEVs from DKs-8 cells under the different conditioning media were also loaded for Western blot analysis based on equal protein concentrations and equal vesicle numbers. The samples were run at 110 V for 1.5 h. Proteins were transferred to nitrocellulose membranes overnight at 15 volts at 4°C. Membranes were stained with Ponceau S solution (cat no. P7170, Sigma) before blocking. Membranes were blocked in 5% non-fat dry milk diluted in Tris-buffered saline with 0.1% Tween 20 (cat no. P9416, Sigma) (TBS-T) for 1 h at room temperature. Primary antibodies were diluted in 5% non-fat dry milk -TBS-T (Ago2, 1:5,000; hnRNP A2/B1, 1:5,000; Syncrip, 1:5,000; YB1, 1:5,000; CD63, 1:10,000; Flotillin, 1:5,000; Hsp70, 1:5,000; Tsg101, 1:5,000; ApoB-100, 1:10,000) and incubated overnight at 4°C. Membranes were washed 3 times for 10 min in TBS-T and subsequently incubated with species-specific HRP-conjugated secondary antibodies (1:10,000; Promega) in 5% non-fat dry milk -TBS-T for 1 h at room temp. All membranes were washed 3 times for 10 min in TBS-T and incubated with an enhanced chemiluminescence (ECL) reagent (cat no. 32106, Thermo Fisher Scientific) or partial amount of femto ECL (cat no. 34095, Thermo Fisher Scientific) for 2 min before being scanned/exposed using an Amersham 680 imager (GE) or an iBright Imager (Thermo Fisher Scientific). Multiple exposures were taken for each blot to have the complete dynamic range for densitometry measurements. The densitometry measurements for the protein bands were done using the Analyze Gels feature of ImageJ (NIH) (Schneider et al., 2012).

### Preparation of Negatively Stained Grids for Transmission Electron Microscopy (TEM)

EV fractions were adhered to freshly glow discharged 300 mesh carbon grids (Electron Microscopy Sciences) by floating the grids on top of 5 µL of samples for 20 seconds. Samples were washed by briefly touching the grid to two, 40 µL drops of ddH_2_O followed by incubation with 5 µL of 2% uranyl acetate for 20 seconds. The negative stain was blotted by pressing the side of the TEM grid against a freshly torn piece of #1 Whatman filter paper until the grid was dry. TEM imaging was performed on a Tecani T12 operating at 100 keV using an AMT nanosprint CMOS camera. Random fields of EVs were imaged over the entire grids.

### Dot blot

As previously described (Lai et al., 2015; McKenzie et al., 2016; Patel & Weaver, 2021; Sung et al., 2015), SEVs were dotted onto nitrocellulose membranes at different protein concentrations (0μg, 0.3μg, 0.6μg, 1.25μg 2.5μg, and 5μg). The dotted membranes were subsequently allowed to dry at room temp for 1 h and then blocked with 5% milk in TBS overnight at 4° C. The membranes were then incubated with anti-rat Ago2 or anti-rabbit CD63 antibody in either 5% milk in TBS or TBS-T and the blots were washed and developed as described in the Western blot section.

### RNase and detergent sensitivity assay

Small EV pellets resuspended in PBS were divided in three equal parts and mixed with buffer, buffer + RNase Cocktail enzyme mix (RNase A + RNase T1) (cat no. AM2286, Thermo Fisher Scientific) (Final concentration RNAse A 5 units and RNase T1 200 Units), or buffer + RNAse Cocktail mix + Triton-X-100 (cat no. T8787, Sigma) (final concentration 1%) in 100 μL and incubated for 30 min at 37°C. Next, 700 μL Qiazol was added, followed immediately by total RNA extraction using the miRNeasy kit (cat no. 217004, Qiagen) according to the manufacturer’s protocol. RNAs were eluted with two rounds of 40 µL of Nuclease-free water (cat no. AM9932, Thermo Fisher Scientific). The quality and size range of each RNA sample was visualized with Bioanalyzer chips, following manufacturer’s protocol; Small RNA kit chips (cat no. 5067-1548, Agilent) with small RNA ladder (4-150nt) were used (cat no. 5067-1550, Agilent), all accessory reagents were also from Agilent (cat no. 5067-1549) using the Agilent 2100 bioanalyzer (G2939A).

### qRT-PCR for small RNAs

Taqman small RNA assays (cat no. 4427975, Thermo Fisher Scientific, U6 snRNA: assay ID 001973; hsa-let7a-5p: assay ID 000377; hsa-miR-100-5p: assay ID 000437; hsa-miR16: assay ID 000391, and hsa-miR-93: assay ID 001090) were performed for RNAs extracted from equal numbers of small EVs according to the manufacturer’s protocol. TaqMan MicroRNA Reverse Transcription Kit (cat no. 4366597, Thermo Fisher Scientific) was used to perform reverse transcription reactions were using RNA extracted from equal number of vesicles in a final reaction volume of 10 μL. After transcription, 0.34 ng (0.67 μL) cDNA was used as the template together with the corresponding Taqman miRNA probe with the TaqMan Universal Master Mix II, no UNG (cat not. 4440049, Thermo Fisher Scientific) for qPCR in a final reaction volume of 10 μL. Each Taqman miRNA qPCR was performed with technical triplicates on a Bio-Rad CFX96. C(t) values were averaged for each technical triplicate. To calculate fold changes (FC), the ΔC(t) method was used (Schmittgen & Livak, 2008). Briefly, ΔC(t) values were calculated for each biological sample, where ΔC(t) = C(t)miRNA. Relative fold changes were determined by Fold change = 2^-ΔΔC(t)^, where ΔΔC(t) = ΔC(t)-ΔC(t)control. For ΔΔC(t) values < 0 (signifying a negative fold change), the negative reciprocal Fold Change formula was used (-1/(2-ΔΔC(t)). Paired t tests were performed from three independent biological replicates.

### Statistics

Most of the experimental data were acquired from at least three independent experiments. The NTA data were compared using Welch’s *t* test and plotted as mean and standard error of the mean in bar graphs in GraphPad Prism 9. For the WB quantitations in Figure 2, paired t tests were used. For the miRNA qRT-PCR data in Figure 3, paired t tests were used. For the WB quantitations in Figure 4 and Figure S5, two-way ANOVA Tukey’s multiple comparisons tests were performed.

**Figure 2.**
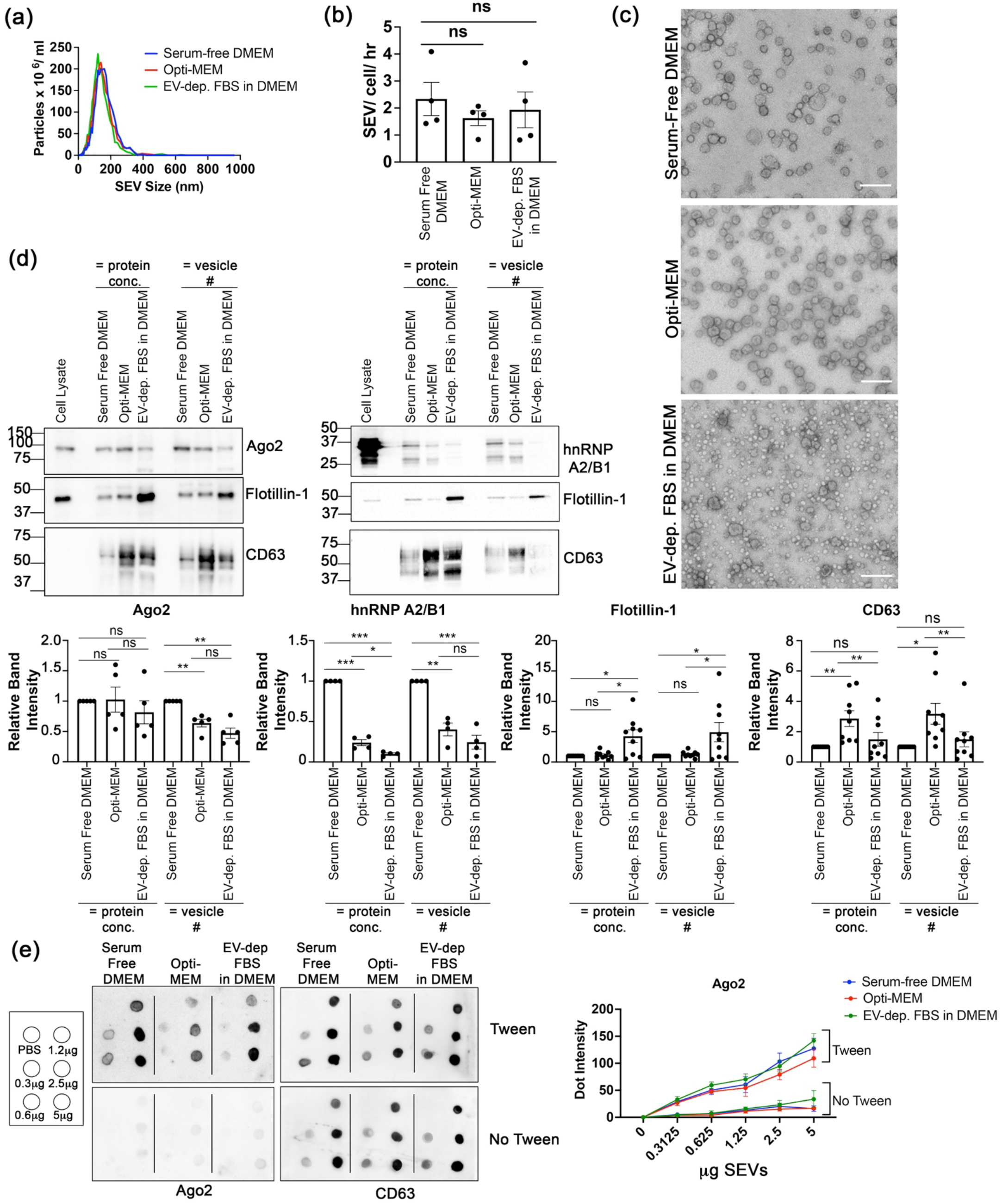
Cell culture media components affect the abundance of Ago2, hnRNP A2/B1, and EV markers in EVs. (a) Representative traces from nanoparticle tracking analysis of DKs-8 SEVs (Fractions 6 and 7) obtained from different conditioning media. (b) Quantitation of DKs-8 SEVs numbers from nanoparticle tracking analysis (n=4). Data were shown as mean + S.E.M. ns, not significant. (c) Representative TEM images of DKs-8 SEVs obtained from the different conditioning media. Scale bar shows 200 nm. (d) Western blot assessments of DKs-8 total cell lysate and SEVs (loading based on equal protein concentration and equal vesicle number) from the different conditioning media, probing for Ago2, hnRNP A2/B1, Flotillin, and CD63 (n=4 or 5). Relative intensity of the Ago2, hnRNP A2/B1, Flotillin and CD63 bands of SEVs in C, normalized to the band intensities of Serum Free DMEM. Paired *t* test, *p=0.05, ***p<0.001, ns, not significant. (e) Different concentrations of SEVs obtained from DKs-8 cells conditioned with the different conditioning media were dotted on nitrocellulose membranes and probed with anti-Ago2 (rat antibody) or anti-CD63 antibodies in the presence (Tween) or absence (No Tween) of 0.1% Tween-20. Right: Quantitation of Ago2 dots in presence or absence of Tween (n=3).

**Figure 3:**
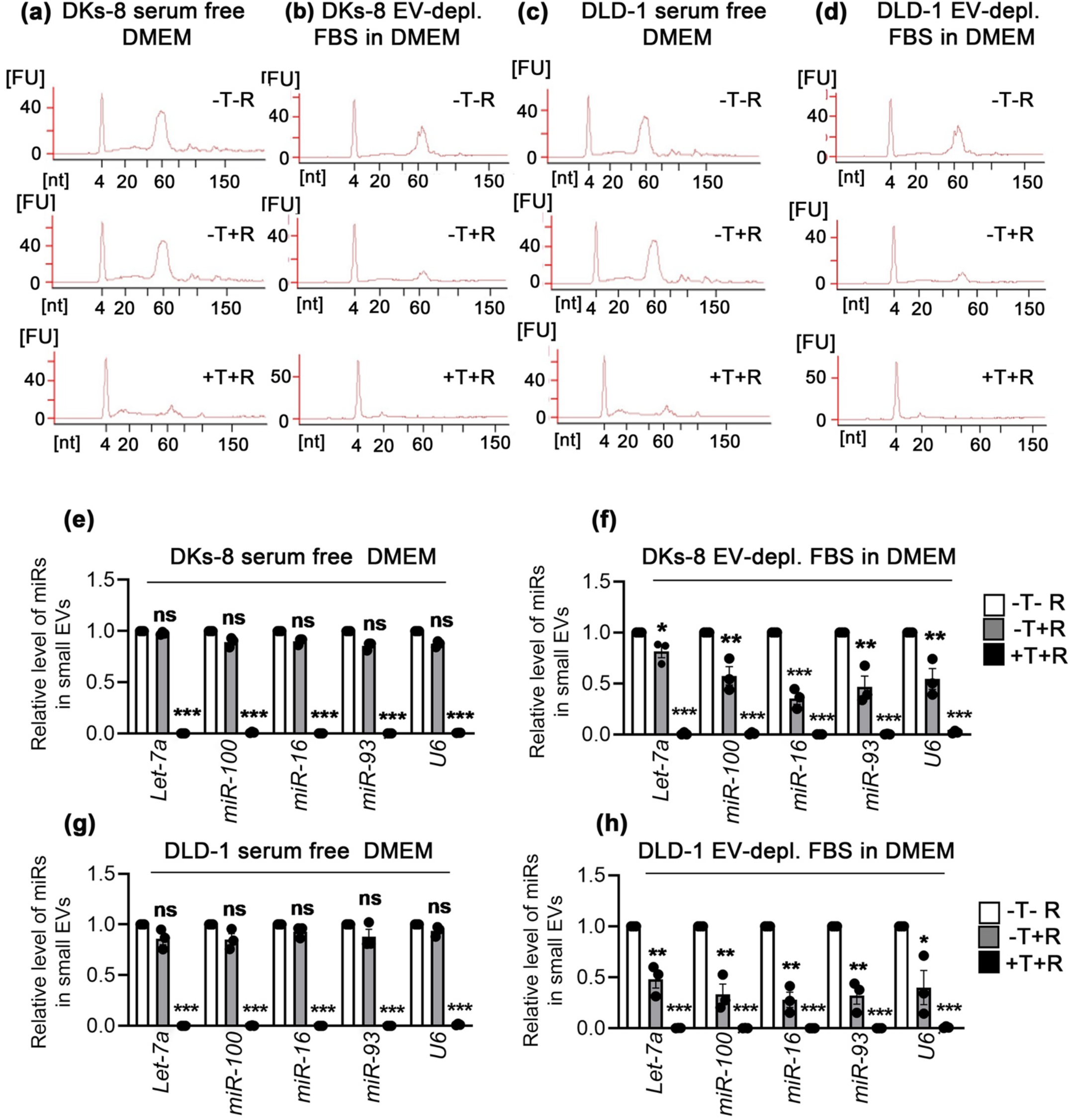
Analysis of small EV associated RNA purified from Serum-free and EV-depleted FBS in DMEM conditioned cells. (a-b) Bioanalyzer electropherogram analysis of purified RNA. Arbitrary fluorescence units (FU) are plotted as a function of RNA size in nucleotides (nt). RNA was extracted from equal numbers of DKs-8 small EVs after detergent (+/- TX100) and RNAse treatment (+/- RNase cocktail). Cells were cultured in serum free DMEM or EV depleted FBS in DMEM conditions. Representative of three experiments, (c-d) Bioanalyzer electropherogram analysis of purified RNA. Arbitrary fluorescence units (FU) are plotted as a function of RNA size in nucleotides (nt). RNA was extracted from equal numbers of DLD-1 small EVs after detergent (+/- TX100) and RNAse treatment (+/- RNase cocktail). Cells were cultured in serum free DMEM or EV depleted FBS in DMEM conditions. Representative of three experiments. (e-f) Relative levels of small RNAs (*let-7a*, *miR-100*, miR-16, *miR-93, and U6*) in small EVs purified from DKs-8 cells cultured in serum-free DMEM or EV-depleted FBS in DMEM conditions. Equal numbers of small EVs were treated with (+) or without (−) RNase cocktail in absence (−) or presence (+) of 1% Triton-X-100 (TX100) followed by total RNA isolation and qRT-PCR. All experiments were done in three biological replicates with three technical replicates. Data plotted as Mean and S.E.M. *p<0.05, **p<0.01, ***p<0.001. (g-h) Relative levels of small RNAs (*let-7a*, *miR-100*, miR-16, *miR-93, and U6*) in small EVs purified from DLD-1 cells cultured in serum-free DMEM or EV-depleted FBS in DMEM conditions. Equal numbers of small EVs were treated with (+) or without (−) RNase cocktail in absence (−) or presence (+) of 1% Triton-X-100 (TX100) followed by total RNA isolation and qRT-PCR. All experiments were done in three biological replicates with three technical replicates. Data plotted as Mean and S.E.M. *p<0.05, **p<0.01, ***p<0.001.

**Figure 4.**
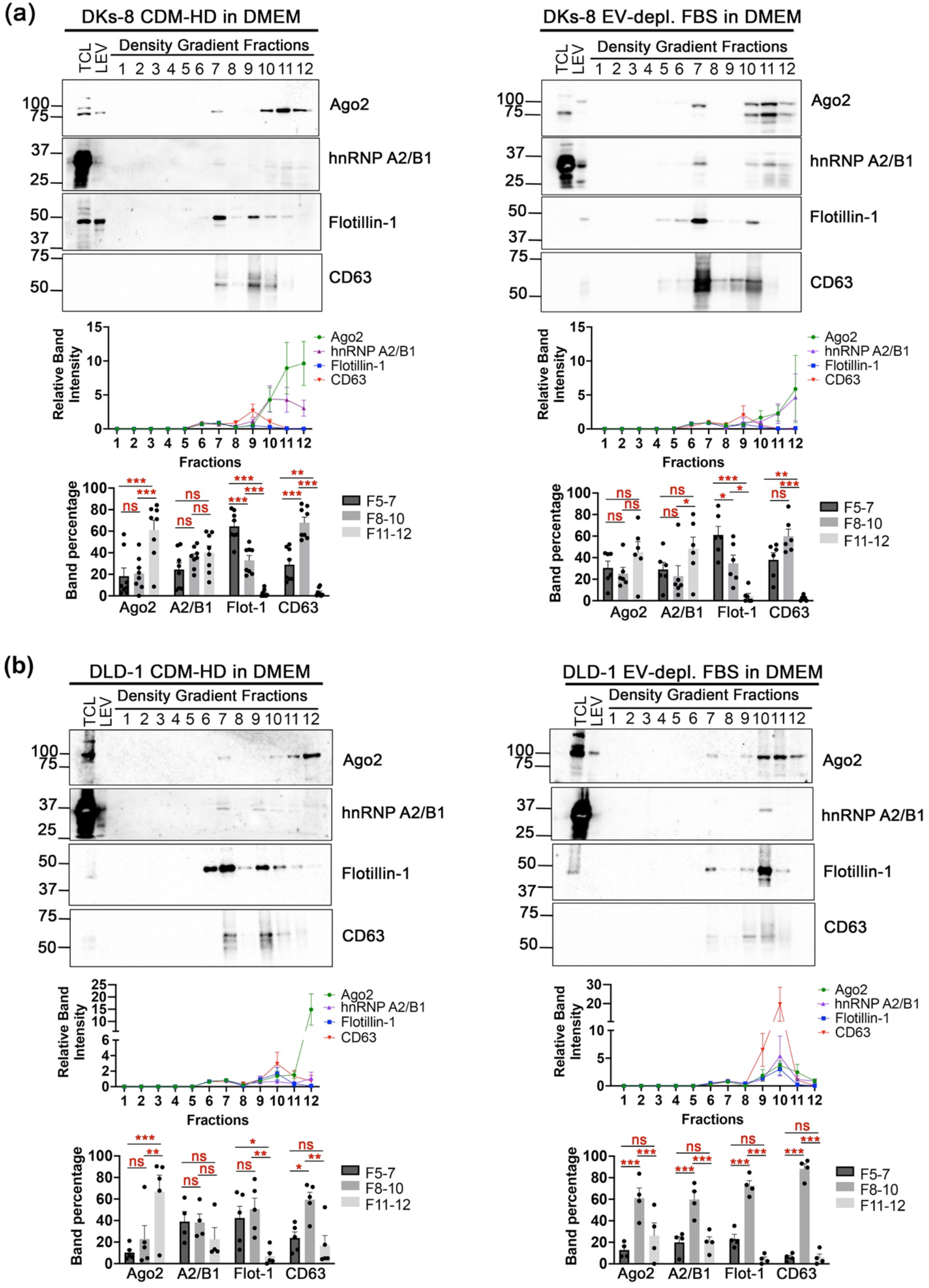
Hollow fiber bioreactors greatly affect EV population and RNA-binding proteins distribution. (a) Western blot analyses of DKs-8 TCL, LEV, and DG fractions from cushion density gradients for CDM-HD in DMEM (Serum-free condition) and EV-depleted FBS in DMEM probed for Ago2, hnRNP A2/B1, Flotillin-1, and CD63. <u>Middle panel:</u> Relative intensity of the Ago2, hnRNP A2/B1, Flotillin-1, and CD63 bands in the cushion density gradient fractions from the different media conditions, normalized to the band intensity of the highest SEV lane (either Fraction 6 or 7) (n=8 and 6, respectively). <u>Bottom panel:</u> Percentage of Ago2, hnRNP A2/B1 (abbreviated A2/B1), Flotillin-1 (abbreviated Flot-1), and CD63 bands found associated with Fractions 5-7, 8-10, and 11-12. Data plotted as Mean and S.E.M. *p<0.05, **p<0.01, ***p<0.001. (b) Western blot analyses of DLD-1 TCL, LEV, and DG fractions from cushion density gradients for the different media conditions probed for Ago2, hnRNP A2/B1, Flotillin-1, and CD63. <u>Middle panel:</u> Relative intensity of the Ago2, hnRNP A2/B1, Flotillin-1, and CD63 bands in the cushion density gradient fractions from the different media conditions, normalized to the band intensity of the highest SEV lane (either Fraction 6 or 7) (n=5 and 4, respectively). <u>Bottom panel:</u> Percentage of Ago2, hnRNP A2/B1 (abbreviated A2/B1), Flotillin-1 (abbreviated Flot-1), and CD63 bands found associated with Fractions 5-7, 8-10, and 11-12. Data plotted as Mean and S.E.M. *p<0.05, **p<0.01, ***p<0.001.

## Results

### Serum is a major contributor of non-vesicular Ago2 in conditioned media

To evaluate the effect of growth factor signaling and serum contamination on the detection of Ago2 in SEVs, we conditioned the colon cancer cell line, DKs-8, with three different culture media (serum-free DMEM, EV-depleted FBS in DMEM, and Opti-MEM) for 48 h. Opti-MEM is a serum-free, but growth factor-replete medium. There was no effect of any of the culture conditions on cell viability (Figure S1a). After removing cell debris and isolating large EVs (LEVs) by differential centrifugation, SEVs were purified from the supernatant utilizing a modified cushion-density gradient protocol (Li et al., 2018). This method eliminates intermediate pelleting steps and potential aggregation of vesicles, with the isolated SEVs collected into a pellet only after the final density gradient fractionation step (Li et al., 2018). We analyzed all fractions across the density gradients, LEVs, and cell lysates by immunoblotting. SEVs were most abundant in density gradient fractions 6 and 7 for all three media conditions, as assessed by detection of the EV marker proteins Flotillin-1 and CD63 (Figure 1a). For the serum-free DMEM and Opti-MEM conditions, Ago2 was primarily detected in the SEV-enriched fractions 6 and 7 (87% and 75%, respectively). By contrast, analysis of density gradient fractions from DMEM supplemented with EV-depleted FBS condition revealed the presence of Ago2 in fractions 6 and 7 (38%) and the non-EV-containing fractions 9-12 (62%). Ponceau stains of the blots further revealed that fractions 9-12 in the EV-depleted FBS condition contain a disproportionate amount of protein compared to other media conditions, including a large band consistent with the size of bovine serum albumen (66 kDa) (Figure 1a, Figure S1b). These data demonstrate that conditioned media containing EV-depleted serum contains both vesicular and non-vesicular Ago2.

To further understand the contribution of serum to non-vesicular Ago2, we carried out cushion density gradient purifications from media that was not exposed to cells (unconditioned media). For EV-depleted serum in DMEM, Flotillin-1 but not CD63 was detected in fractions 5-7, suggesting the possible presence of residual EVs. Consistent with previous reports of non-vesicular Ago2 in blood (Arroyo et al., 2011; Geekiyanage et al., 2020), Ago2, but not Flotillin-1 or CD63, was detected only in fractions 9-12 (Figure S1c). The RBP hnRNP A2/B1 was also present in nonvesicular fractions from unconditioned EV-depleted serum in DMEM media (Figure S1c). For unconditioned Opti-MEM media (Figure S1c), Ago2 was undetectable in any of the fractions; however, minor bands for Flotillin-1 and hnRNP A2/B1 were detected in fraction 7, suggesting that this proprietary serum-free media contains low numbers of contaminating EVs and RBPs.

We also evaluated the effects of serum in conditioned media obtained from the mutant KRAS colon cancer cell line, DLD-1. DKs-8 cells are derived from this cell line, having the mutant KRAS gene edited out (Shirasawa et al., 1993); thus, alterations in EV character and content between the two cell lines should be attributable to differences in KRAS signaling. Indeed, we previously found that KRAS mutation downregulates Ago2 trafficking into EVs (McKenzie et al., 2016). Consistent with the distribution of Ago2 from DKs-8 cell culture, conditioning DLD-1 cells in EV-depleted serum-containing media resulted in the detection of Ago2 in both vesicular and non-vesicular fractions. Thus, for both DKs-8 and DLD-1 cell lines, greater than 50% of the isolated Ago2 was associated with nonvesicular fractions 9-12 when cells were conditioned in EV-depleted serum-containing media (Figure 1b). Notably, when DLD-1 cells were conditioned in serum-free media, extracellular Ago2 remained distributed between both vesicular and non-vesicular fractions, a marked departure from the predominantly EV-associated Ago2 isolated from DKs-8 cells cultured under serum-free conditions. While ∼87% of the Ago2 from DKs-8 serum-free conditioned media cofractionates with EVs, only ∼53% of the Ago2 from DLD-1 cultures cofractionates with EVs under the same conditions (Figure 1b). Thus, increased secretion of non-vesicular Ago2 was observed from cells with altered KRAS signaling when cultured under serum-free conditions.

In addition to media additives, alternate purification methods can lead to differential segregation of vesicular and non-vesicular components. A method that has been shown to separate such components with high resolution is bottom-up flotation of EVs from an initial crude EV pellet (Jeppesen et al., 2019; Zhang et al., 2023). To compare our cushion density gradient method with flotation from an EV pellet, we ultracentrifuged conditioned media gathered from DKs8 cells cultured in EV-depleted serum in DMEM for 4 h at 120,000 x g. The crude EV pellet from that centrifugation was then resuspended and placed at the bottom of a 5-40% iodixanol density gradient and centrifuged overnight at 100,000 x. g. Similar to our results obtained by cushion density gradient isolation of SEVs (Figure 1a), fractions isolated using a 4-hour ultracentrifugation-to-density gradient protocol contained Ago2 in both SEV (CD63 and flotillin-containing) fractions and non-vesicular fractions (Figure 1c). These data are consistent with the packaging and secretion of Ago2 in SEVs by DKs-8 cells as well as a contribution of nonvesicular Ago2 from the EV-depleted serum (Figure S1C).

### Serum-free conditions enhance the release of SEVs enriched in RBPs

To quantify the number and size of SEVs obtained under different cell culture conditions, we combined the cushion density gradient purified SEVs from peak fractions 6 and 7 and performed nanoparticle tracking analysis. No significant differences were observed in the number or size of the SEVs collected using the three conditioning methods (Figures 2a and 2b). Transmission electron microscopy (TEM) imaging of negatively stained samples further demonstrated that the EVs collected from serum-free and Opti-MEM media conditions were relatively uniform with sizes of approximately 100 nm (Figure 2c). EVs of similar character were also observed in TEM images of SEVs purified from EV-depleted FBS in DMEM; however, copious numbers of very small vesicular structures were also visible (Figure 2c). Based on their size of ∼25 nm, we hypothesized that these may be low density lipoprotein (LDL) particles. Indeed, Western blot analysis of SEVs from the DKs-8 EV-depleted FBS in DMEM condition revealed a significant level of co-isolating Apolipoprotein B-100, the major protein component of LDL (Figure S2a).

Immunoblots of the pooled peak fractions were then carried out to provide direct side-by-side comparisons of Ago2 levels in the SEVs secreted from DKs-8 cells under the different media conditions. SEVs were loaded for Western blot analysis based on either equal protein concentrations or equal vesicle numbers. When loaded based on equal protein levels, Ago2 levels were not significantly altered by the different media conditions (Figure 2d, Figure S2b). However, loading based on equal vesicle number revealed that Ago2 levels were significantly reduced in SEVs obtained from both the Opti-MEM and EV-depleted FBS in DMEM conditions compared to the Serum-free DMEM condition (Figure 2d). Flotillin-1 levels were significantly upregulated in SEVs from the EV-depleted FBS in DMEM condition, which may be due to flotillin-positive lipid entities in the serum (Figure 2d, Figure S1c). Conversely, conditioning cells with Opti-MEM consistently generated SEVs with significantly higher CD63 content (Figure 2d), presumably due to signaling changes induced by undisclosed components of this proprietary formulation. To determine whether the impact of media composition on Ago2 content might portend an effect on other RBP cargo, we also examined the levels of another EV-associated RNA-binding protein, hnRNP A2/B1. Indeed, hnRNP A2/B1 levels were significantly reduced in the SEVs derived from Opti-MEM and EV-depleted FBS in DMEM conditions compared to Serum-free DMEM for both gel loading conditions (equal protein or equal vesicle, Figure 2e). Taken together, these studies indicate that conditioning DKs-8 cells in serum-free DMEM is superior to the other tested media for production of SEVs low in lipoprotein contamination and high in RBP cargo.

As both EV-depleted serum and Opti-MEM contain growth factors, we next sought to determine whether growth factor signaling might account for the decreased Ago2 present in SEV preparations from those media conditions. To test this hypothesis, we supplemented serum-free DMEM with epidermal growth factor (EGF, 100 ng/mL) (Larios et al., 2020; Oszvald et al., 2020) and immunoblotted for Ago2 in the SEVs secreted under this condition compared to standard serum-free DMEM. No significant change in SEVs numbers were observed under the two conditioning media (Figure S3a). Likewise, Ago2 levels in the isolated SEVs were unchanged by addition of EGF (Figure S3b). Thus, EGF content in conditioning media is not the key factor regulating Ago2 trafficking to SEVs.

One approach used to differentiate bona fide SEV cargoes from co-isolating contaminants is to analyze whether the protein can be detected in the absence of membrane permeabilization. Prior studies by our laboratory and others have reported Ago2 to be detected only in the presence of detergent, i.e., localized inside the SEV membrane (Mantel et al., 2016; McKenzie et al., 2016), while other studies found that Ago2 could be detected in their SEV preparations without detergent permeabilization (Zand Karimi et al., 2022). To evaluate whether media composition alters the intraluminal presence of Ago2 in highly purified SEVs, we performed a detergent sensitivity assay in a dot blot format (Lai et al., 2015; McKenzie et al., 2016). DKs-8-derived SEVs purified from serum-free DMEM, EV-depleted FBS in DMEM, and Opti-MEM conditioned media, were serially diluted and dotted onto nitrocellulose membranes, and probed for Ago2 or CD63 in the presence or absence of 0.1% Tween-20 detergent. For all media conditions, the vast majority of Ago2 signal was detected only in the presence of Tween-20 (Figure 2e). The extracellular domain of CD63 served as a positive control and indeed was detected in both the presence or absence of detergent (Figure 2e). These data indicate that our cushion density gradient purification method is capable of separating nonvesicular Ago2 from Ago2 that is incorporated into EVs, such that even the EVs purified from serum-containing media had very little detectable extravesicular Ago2.

### EV-depleted FBS is a source of extravesicular RNA

Serum is a known source of non-vesicular RNA in conditioned media (Driedonks et al., 2019; Kornilov et al., 2018; Mannerstrom et al., 2019; Shelke et al., 2014; Wei et al., 2016). Using an RNase and detergent sensitivity assay, we analyzed whether these serum-derived RNA contaminants carry through our cushion density gradient SEV purification. EVs were purified from DKs-8 and DLD-1 conditioned media of either serum-free or EV-depleted serum-containing DMEM and subjected to RNase treatment in the presence or absence of 1% Triton X-100 detergent before the extraction of RNA. Electropherograms of the recovered RNA contained a major peak around 60 nt that disappeared completely in the presence of both Triton X-100 and RNase (Figure 3a-d, bottom graphs). The RNA peak from SEVs isolated from either cell line in serum-free conditioned media remained largely intact with RNase treatment in the absence of detergent. However, when EV-depleted serum was included in conditioning media, the RNA associated with SEVs from either cell line was significantly reduced by treatment with RNAse alone, (Figure 3a-d, middle graphs). We next determined RNAse sensitivity at the level of individual RNAs, analyzing two miRNAs previously identified as cargoes of DKs-8 and DLD-1 SEVs (let-7a and miR-100; (Barman et al., 2022; McKenzie et al., 2016), two miRNAs previously reported to be serum contaminants (miR-16 and miR-93; (Mannerstrom et al., 2019), and U6 snRNA which is frequently used as an SEV cargo positive control. For SEVs secreted under serum-free culturing conditions, all four miRNAs and U6 were unaffected by RNase treatment in the absence of detergent and fully depleted by RNase in the presence of detergent (Figure 3e, g). However, for SEVs purified from EV-depleted serum conditions, RNase treatment in the absence of detergent significantly depleted the levels of all four miRNAs and U6, indicating that these SEV preparations retained significant amounts of nonvesicular small RNAs (Figure 3f, h), despite minimal contamination with extravesicular Ago2 (Figure 2e).

### Hollow fiber bioreactor cell culture greatly affects EV populations and RNA-binding protein distribution

In addition to media components, cell density and the dimensionality of the culturing environment can have major impacts on cellular metabolism and EV biogenesis. To investigate the contribution of these factors, we expanded our assessment to EVs obtained from cells cultured in commercial hollow fiber bioreactors. DKs-8 and DLD-1 cells were grown in bioreactors under two media conditions that support high density cell culture: serum-free DMEM supplemented with CDM-HD, a chemically defined, protein-free growth supplement obtained from the bioreactor company (FiberCell Systems), or EV-depleted FBS containing DMEM media. Conditioned media was collected for either 24 (DLD-1 cells) or 48 hours (DKs-8 cells) and subjected to the cushion-density gradient protocol to isolate the SEV-containing fractions. in contrast to the uniform population of SEVs isolated in fractions 6+7 from 2D culture conditions, immunoblotting and nanoparticle tracking analysis revealed that SEVs from hollow fiber bioreactors fractionated into two distinct peaks. Detection of SEV marker proteins Flotillin-1 and CD63 peaked in fractions 6 and 7 and again in fractions 9 and 10, correlating with high particle counts in both populations (Figure 4a and Figure S4). Consistent with these observations, transmission EM images of particles purified from pooled fractions 6+7, 9+10, and 12 revealed SEVs in both 6+7 and 9+10 pooled fraction peaks, with only nonvesicular protein aggregates visible in fraction 12 (Figure S4). Notably, fractions 9+10 exhibited higher levels of nonvesicular protein (visualized as the darkly stained material across grids) than fractions 6+7. Whether this reflects proteins bound to EVs or unassociated protein that co-isolates with the EVs is unclear.

In contrast to 2D culture of DKs-8 cells (Figure 1), Ago2 and hnRNP A2/B1 were associated with both vesicular and non-vesicular fractions isolated from hollow fiber bioreactors irrespective of media composition (Figure 4a). Moreover, the majority of Ago2 and hnRNP A2/B1 appeared to be non-vesicular, resolving to fractions 11 and 12, (Figure 4a). We additionally assessed the presence of other previously reported EV-associated RBPs, such as YB1 and SYNCRIP (Barman et al., 2022; Kossinova et al., 2017; Santangelo et al., 2016). These RBPs were also largely found in the late density gradient fractions containing nonvesicular protein (Figure S5).

Analysis of the conditioned media obtained from DLD-1 cells cultured in hollow fiber bioreactors revealed distinct differences in the vesicular and non-vesicular association of RBPs compared to DKs-8 cells. Like the fractions of the DKs-8 conditioned media from the hollow fiber bioreactors, Ago2 was found in both vesicular and non-vesicular fractions from the EV-depleted FBS condition of the DLD-1 cells (Figure 4b). In contrast, the majority of Ago2 was non-vesicular and found in fractions 11 and 12 in the serum-free condition (Figure 4b). This also differed from what we observed with the 2D culture of the DLD-1 cells, where serum-free conditions led to enhanced detection of EV-associated Ago2 (Figure 1b). Immunoblotting of hnRNP A2/B1 revealed that is was present at low levels in conditioned media fractions, with the majority dected in EV-associated fractions from either serum-free or serum-containing conditions (Figure 4b). Immunoblotting of YB1 and SYNCRIP revealed a similar pattern to that of Ago2, with enhanced detection in nonvesicular fractions in the serum-free condition and significant amounts in vesicular fractions (6+7 and 9+10) in the serum-containing media (Figure S5b). These data suggest that cell density and dimensionality have a major impact on the release and detection of vesicular and nonvesicular RBP cargoes.

Hollow fiber bioreactors incorporate a membrane with pores that should exclude proteins >20 kDa. If the protein is spherical, the estimated radius of a 20 kDa protein is ∼ 1.78 nm (Erickson, 2009), suggesting that EVs, lipoproteins, and large proteins should be excluded. The nonvesicular RBPs observed in material collected from bioreactor-conditioned media should thus be primarily derived from the cells and not from media components. To confirm that proteins and vesicles from media do not cross this membrane, we analyzed unconditioned EV-depleted FBS in DMEM media collected from a bioreactor that had not been inoculated with any cells. Neither EV markers nor RBPs were detected by immunoblot in this material (Figure S6a). We also did not detect ApoB-100, indicating no contamination with LDL (Figure S6b). Ponceau staining of the blots revealed that there was a small amount of protein derived from the media, notably a band at the approximate size of bovine serum albumen (66 kDa); however, the amount was greatly reduced from that observed in cushion density gradients of the same media from 2D culture (compare Figures S1c and S6b). TEM analysis of the fractions further confirmed that very few SEVs and protein aggregates were present (Figure S6c). Thus, most but not all EVs and proteins are excluded by the bioreactor membrane.

## Discussion

In this study we set out to determine how cell culture conditions can affect the association of RBPs and RNAs with SEVs. We found that in 2D culture conditions, serum contributes a substantial amount of both RBPs and RNA to the conditioned media. Furthermore, both nonvesicular RNA and lipoproteins were detected in highly purified EV preparations from EV-depleted serum-containing media. In addition, serum-containing and Opti-MEM media also diminished the levels of intravesicular Ago2 and hnRNP A2/B1, compared to serum-free DMEM. Finally, examination of the impact of culturing cells in a commonly used commercial bioreactor revealed that a large amount of nonvesicular RBPs is present in conditioned media, especially under serum-free conditions. Altogether, these data indicate that the culture conditions greatly influence the RBP and RNA levels both inside and outside of EVs and should be carefully considered for EV RNA studies.

Serum-containing media contains a wide variety of proteins, lipoproteins, nonvesicular RNA-RBP complexes and EVs and has been previously reported to interfere with interpretation of EV analyses (Arroyo et al., 2011; Lehrich et al., 2018; Mannerstrom et al., 2019; Tosar et al., 2017; Turchinovich et al., 2011; Wei et al., 2016). One strategy that some investigators use is to include EV-depleted FBS in their conditioning media, although it has been reported that an 18h ultracentrifugation of FBS does not completely deplete the FBS of EVs, with about 30% of FBS-EVs remaining (Lehrich et al., 2018). Furthermore, it leaves behind a large amount of nonvesicular RNA-RBP complexes and lipoproteins, as we found in our study. Consistent with our finding that LDL from EV-depleted serum copurified with EVs upon flotation in density gradients (Figure 2 and Figure S2), contaminating LDL was previously detected in EV-depleted FBS (Gardiner et al., 2013). High-density lipoproteins (HDL) are another potential contaminant in density gradient preparations (Brennan et al., 2020). Notably, like EVs, lipoproteins are carriers of extracellular RNAs and could be additional sources of non-EV based RNAs (Vickers et al., 2011) beyond nonvesicular RNA-RBP complexes. A final consideration is that supplements added to serum-free media may also contain miRNAs and may be co-isolated in EV collection (Auber et al., 2019).

The incorporation of cargoes into EVs can be impacted by many factors, including the signaling and metabolic state of the cells (Dixson et al., 2023). For colon cancer, a major driver of oncogenic signaling is KRAS mutation. Demory Beckler *et al*. reported that KRAS mutation in isogenic colon cancer cells greatly affects the protein composition of SEVs, with a greater proportion of RBPs present in SEVs purified from wild-type colon cancer cells (Demory Beckler et al., 2013). Consistent with the model in which RBPs carry RNA into EVs (Dixson et al., 2023; O’Brien et al., 2020), Cha *et al*. reported that specific non-coding RNAs, such as miRNAs, rRNAs, tRNAs and scRNAs, are differentially sorted to SEVs depending on the KRAS status of colon cancer cells (Cha et al., 2015). We further previously reported that sorting of Argonaute 2 (Ago2) and several miRNAs to exosomes/SEVs is controlled by KRAS-MEK-ERK signaling in colon cancer cells (McKenzie et al., 2016). Specifically, phosphorylation of Ago2 on serine 387 prevents Ago2 association with multivesicular endosomes (MVE) and sorting into exosomes. Here, we compared wild-type and mutant KRAS responses to the various cell culture conditions. While most responses to culture conditions were similar (e.g. 2 EV peaks and increased nonvesicular RBPs in density gradient preparations of media from hollow fiber bioreactors and contamination of EV preparations from EV-depleted serum media from 2D flasks with nonvesicular RNA), there were a few instances in which mutant KRAS cells had distinct differences. These differences include the presence of substantial amounts of nonvesicular Ago2 in serum-free 2D conditions in mutant but not wild-type KRAS CRC cells (Figure 1) and the presence of nonvesicular YB1 in serum-containing bioreactor conditions for wild-type but not mutant KRAS cells (Figure S5). Overall, however, the culture conditions seemed to be a more dominating factor than KRAS mutation status.

Bioreactors are a convenient way to scale up cell cultures and obtain enough media for large-scale EV purifications for proteomics, RNA-Seq, animal experiments, or commercial purposes. Here we compared the preparations from a commonly used commercial hollow fiber bioreactor (FiberCell) to those we obtained from 2D flask culture. There were several differences between bioreactor and 2D flask culture. First, using identical flotation density gradient protocols, there were two distinct EV peaks coming from bioreactor culture media as compared to a single EV peak from flask culture. These peaks are likely biologically distinct, as Flotillin tended to be enriched in the fraction 6+7 peak (except for DLD-1 cells in serum-containing media) whereas CD63 tended to be enriched in the fraction 9+10 peak (Figure 4). An interesting future direction could be to see if there are additional differences in EV cargoes between the peaks. Second, there was a large amount of nonvesicular RBPs present in media collected from bioreactor cultures (fraction 12, Figure 4 and Figure S5) that was not derived from the media (Figure S6). The origin of these nonvesicular particles is unknown but could be exported in a specific manner from cells (Zhang et al., 2021) or released from dying cells in the bioreactor space.

When comparing serum-containing DMEM to serum-free DMEM, we also assessed whether a growth factor-enriched serum-free media Opti-MEM or the addition of EGF to DMEM would impact the levels of RBPs in EVs. Based on our previous work showing that KRAS-MEK-ERK signaling downstream of KRAS mutation downregulates Ago2 trafficking to SEVs (McKenzie et al., 2016), we expected that growth factor signaling would downregulate Ago2 in SEVs. Indeed, for the Opti-MEM condition there was a downregulation of Ago2 in EVs purified from wild type KRAS cells; however, EGF alone was not sufficient to alter Ago2 levels in EVs. As Opti-MEM is a proprietary media, it is unclear what the responsible factor is. Future investigations should further explore the specific cell signaling pathways that regulate trafficking of Ago2 and other RBPs into EVs and how these are impacted by cell media formulations, cell signaling state, and tissue environment.

RNA and RBPs have been reported to be minor EV components (Albanese et al., 2021; Jeppesen et al., 2019). We recently identified a subpopulation of EVs that is highly enriched in RNA and RBPs and is packaged at ER membrane contact sites (Barman et al., 2022). This EV subpopulation was characterized from DKs-8 cells using serum free conditions and represented ∼10% of the EV population (Barman et al., 2022). Given the relative rarity of RNA-containing EVs in a bulk EV population and the confounding factors of cell signaling state and of nonvesicular RNA-RBP complexes in serum and high density culture supernatants, it is not surprising that investigators have had difficulty characterizing RNA and RBPs as bona fide EV cargoes. This current study parses many of the factors that affect both the true signal and the background noise in characterization of RBPs and RNAs in EVs. Consideration of these factors and performing detergent sensitivity analysis of intra- vs. extra-vesicular RNA and RBPs for any given study is highly recommended.

## Author contributions

Conceptualization- L.J and A.M.W.

Data curation- L.J., B.B., Y.J.J., L.C., E.K., C.S.

Formal Analysis- L.J., B.B., Y.J.J.

Funding Acquisition- L.J., J.W., A.M.W.

Investigation- L.J., B.B., Y.J.J., L.C., E.K., C.S.

Methodology- L.J., B.B., Y.J.J., L.C., E.K.

Project Administration- T.R.D., A.M.W.

Resources- A.M.W., K.V.

Supervision- A.M.W., J.W.

Visualization- L.J., B.B., Y.J.J.

Writing-original draft- L.J.

Writing-review and editing- L.J., B.B., Y.J.J., L.C., E.K., C.S., K.V., J.W., T.R.D., A.M.W.

## Acknowledgements

We thank members of the Weaver laboratory and P01 research group for their feedback on this project. We thank Drs. Jeffrey Franklin and Robert Coffey for their assistance with the hollow fiber bioreactors. This work was supported by funding provided by NIH grants F32 CA217064 and P01 CA229123, as well as NSF grant NSF-2036809. Transmission electron microscopy was performed in part through the use of the Vanderbilt Cell Imaging Shared Resource (supported by NIH grants CA68485, DK20593, DK58404, DK59637 and EY08126).

## Conflict of Interest

The authors have declared no conflict of interest.

## Figure Legends

**Figure S1.**
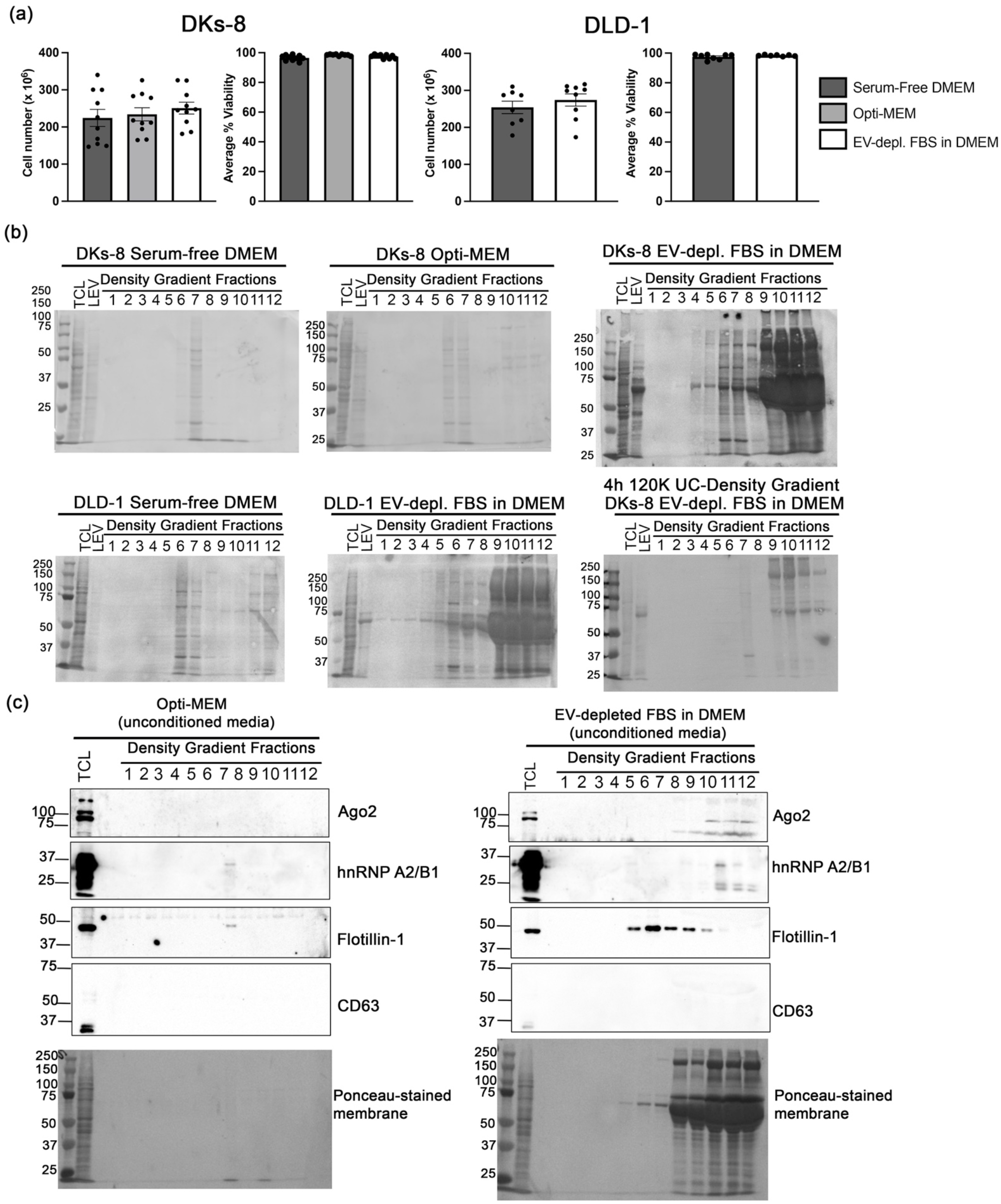
Supporting data for Figure 1. (a) Viability numbers of the DKs-8 and DLD-1 cells at the time of conditioned media collection (n=8-10). (b) Ponceau stained membranes from blots shown in Figure 1. (c) Representative Western blots of overnight cushion density gradient fractions of unconditioned Opti-MEM and EV-depleted FBS in DMEM, probed for Ago2, hnRNP A2/B1, Flotillin-1, and CD63. Ponceau stained membranes of the blots are shown below.

**Figure S2.**
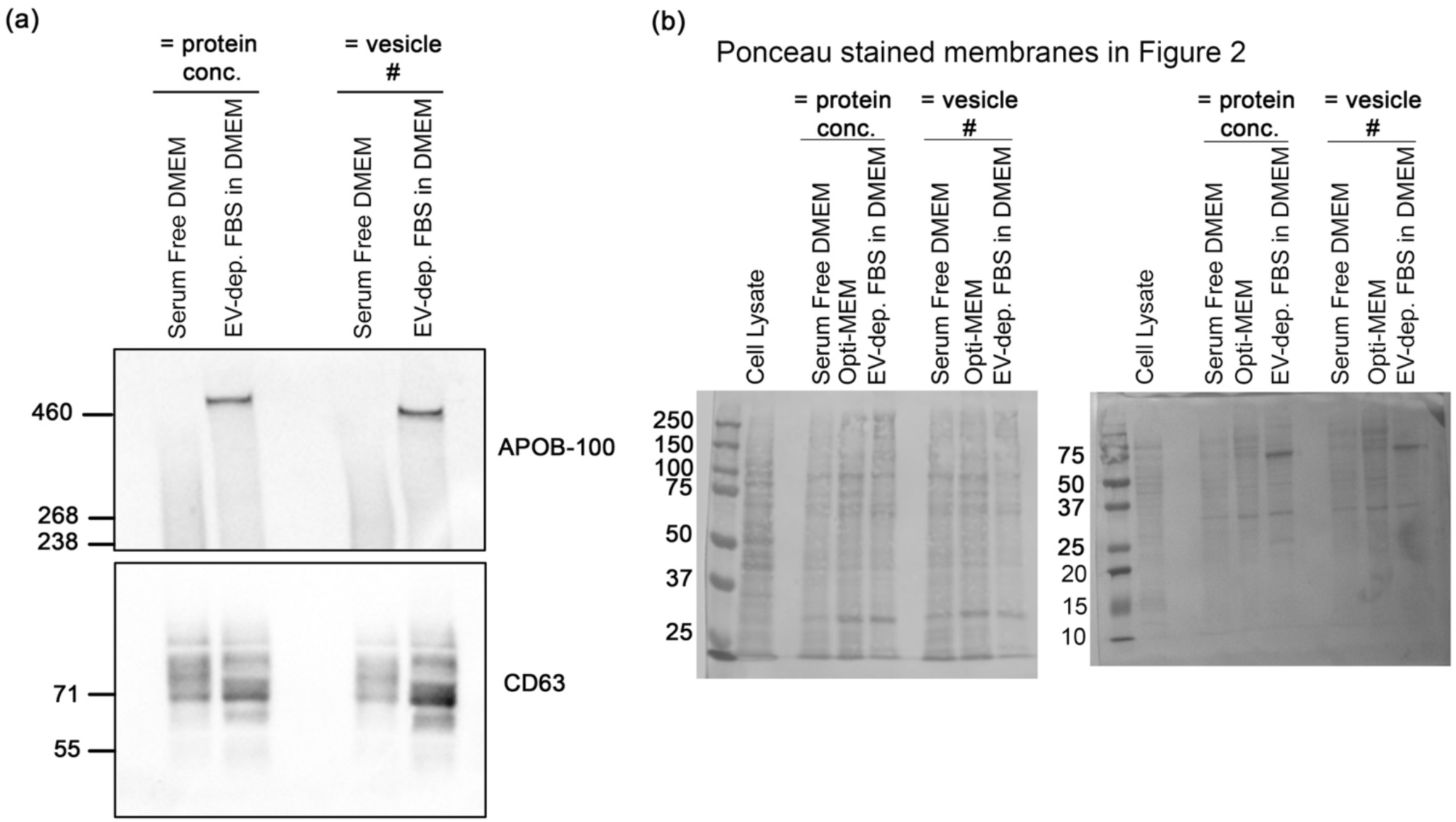
Supporting data for Figure 2. (a) Western blot assessments of DKs-8 SEVs (loading based on equal protein concentration and equal vesicle number) from the Serum-free DMEM and EV-depleted FBS in DMEM conditions, probing for APOB-100 and CD63 (n=3). (b) Ponceau stained membranes from blots shown in Figure 2.

**Figure S3.**
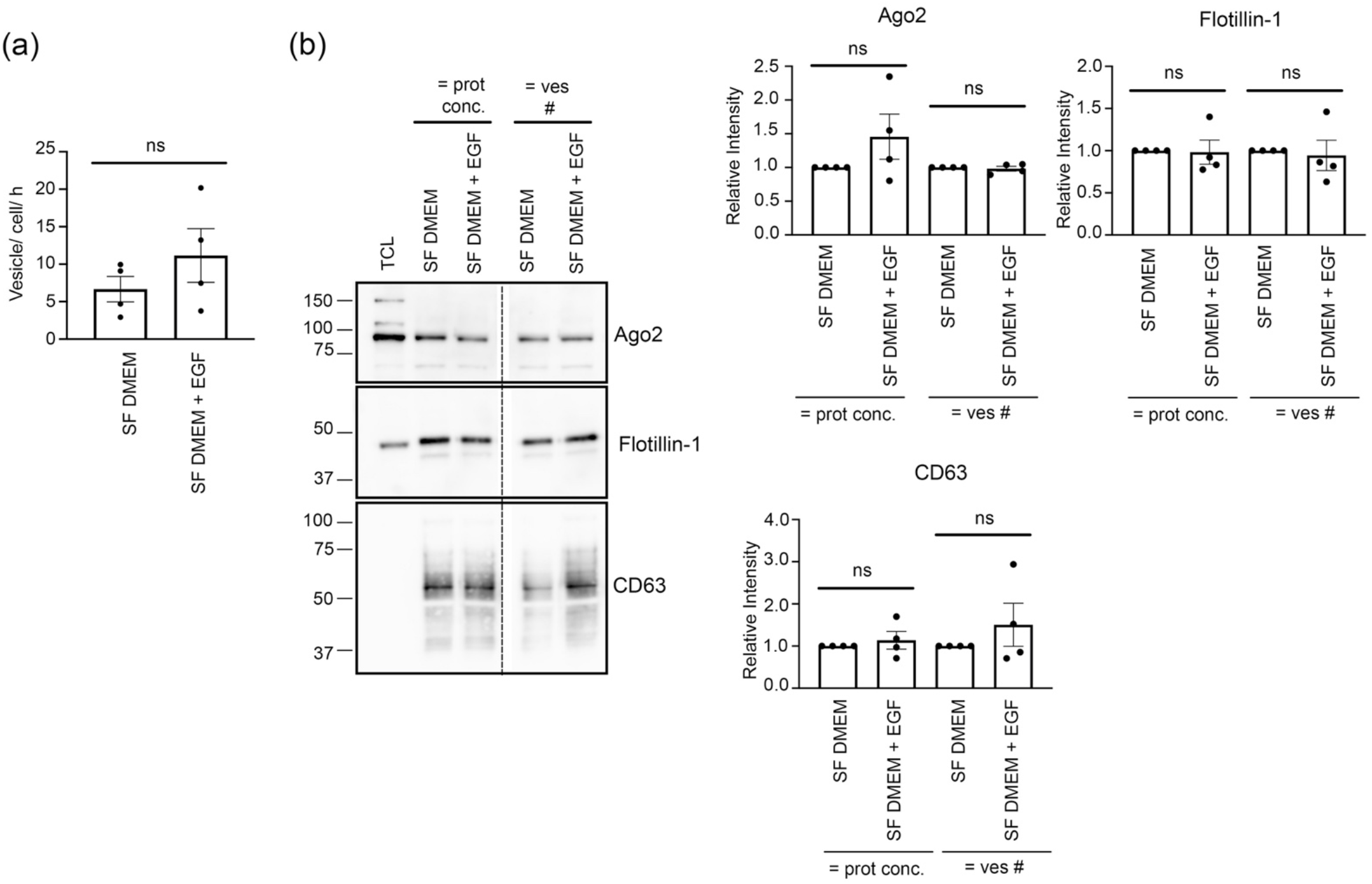
Ago2 levels in SEVs not altered by EGF addition. (a) Quantitation of DKs-8 SEVs numbers obtained from different conditioning media (Serum-Free DMEM, Serum-Free DMEM supplemented with 100 ng/mL EGF, and Opti-MEM) (n=4). Data were shown as mean + S.E.M. ns, not significant. (b) Western blot assessments of DKs-8 total cell lysate and SEVs (loading based on equal protein concentration and equal vesicle number) from the different conditioning media, probing for Ago2, Flotillin, and CD63 (n=4). <u>Right:</u> Relative intensity of the Ago2, Flotillin and CD63 bands of SEVs in C, normalized to the band intensities of Serum Free DMEM. Z-test *p<0.05, ***p<0.001, ns, not significant.

**Figure S4.**
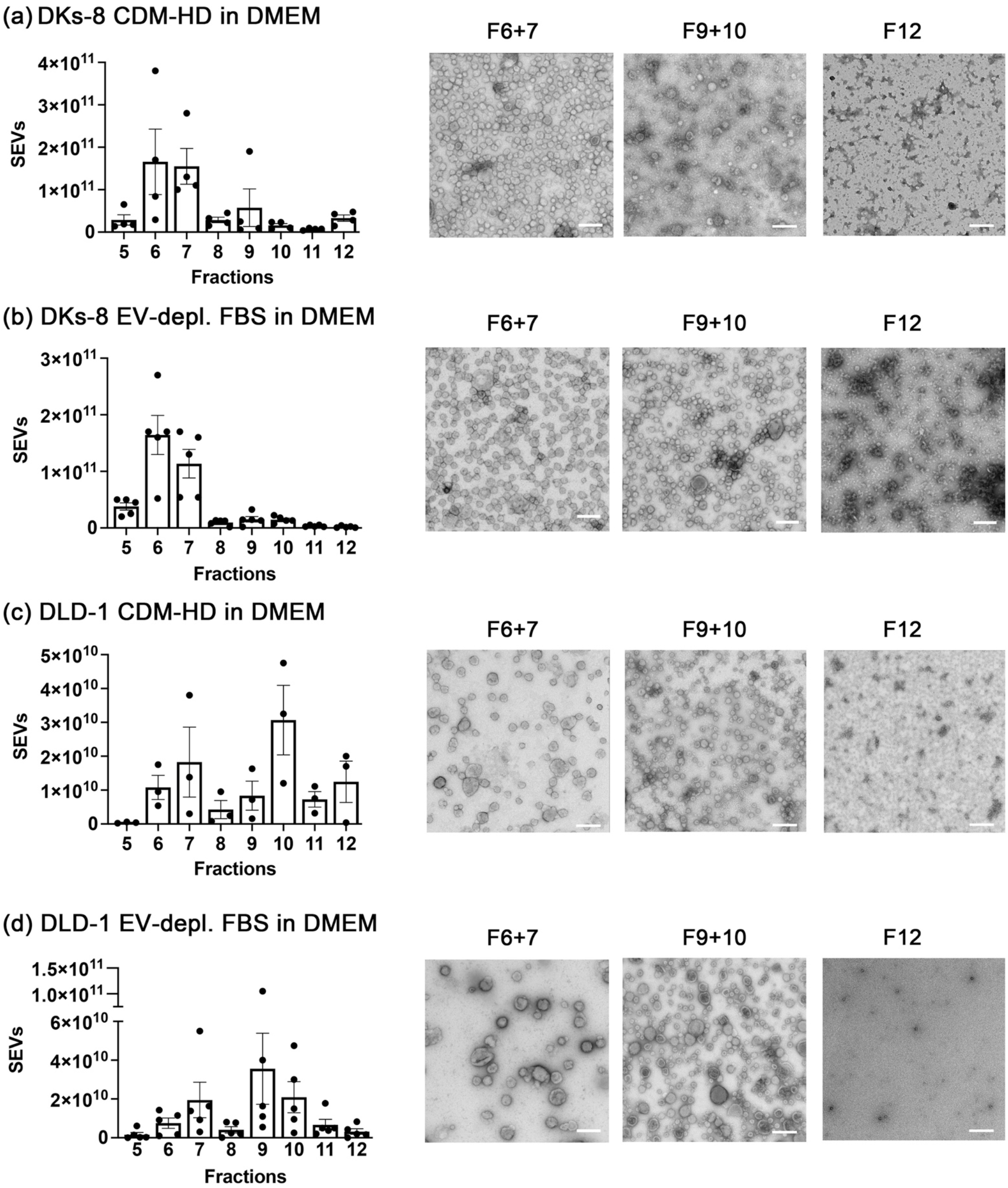
Characterization of density gradient fractions of media from bioreactors. (a-d) SEV numbers in DKs-8 or DLD-1 DG fractions (F5-12) from cushion density gradients for CDM-HD in DMEM (Serum-free condition) and EV-depleted FBS in DMEM determined by nanoparticle tracking analysis (n=3-5). Data plotted as Mean ± S.E.M. <u>Right:</u> Representative TEM images of SEVs obtained from the different conditioning media. Scale bar shows 200 nm.

**Figure S5.**
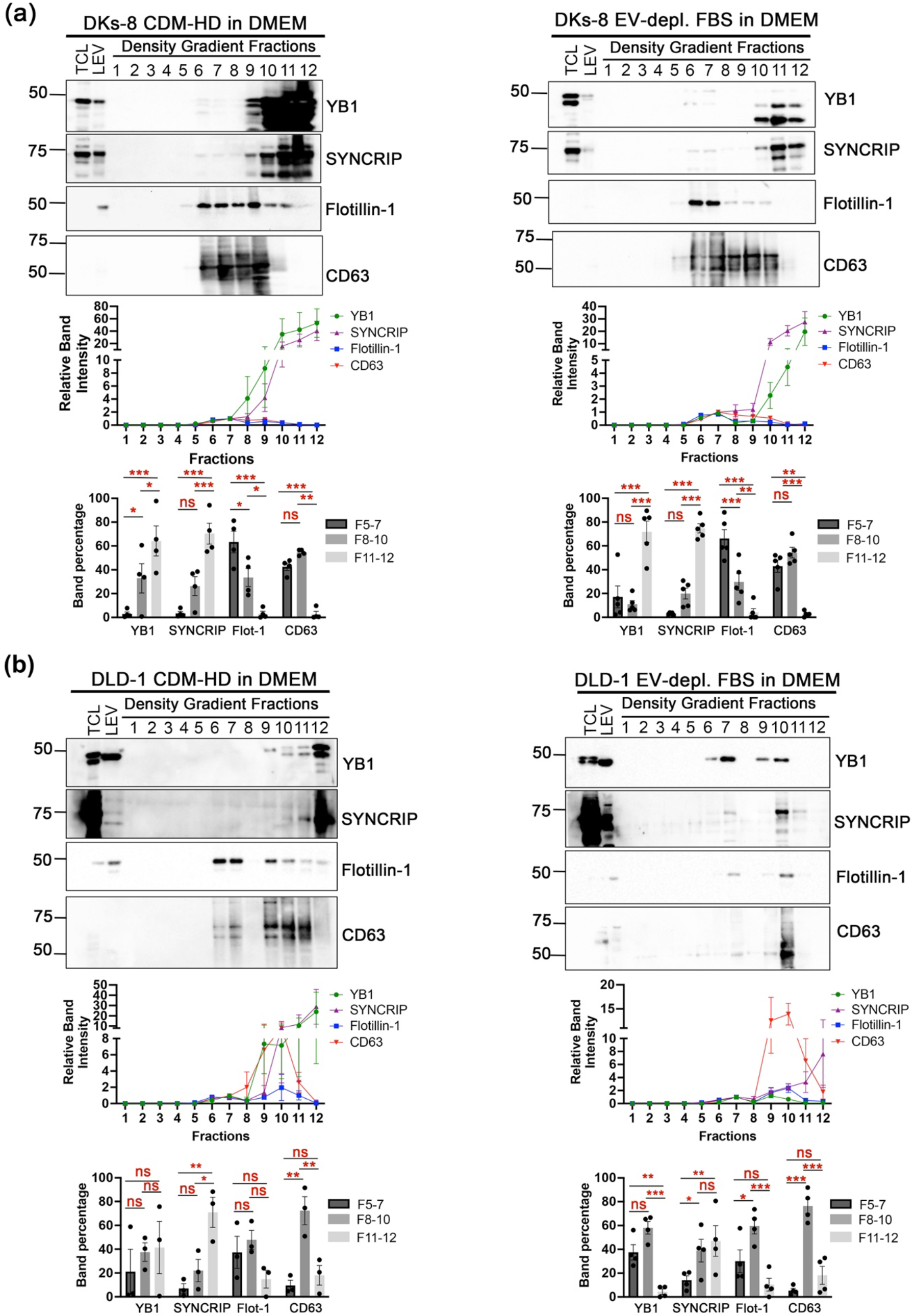
Western blot assessments of additional RNA-binding proteins in density gradient fractions of media from bioreactors. (a) Western blot analyses of DKs-8 TCL, LEV, and DG fractions from cushion density gradients for CDM-HD in DMEM (Serum-free condition) and EV-depleted FBS in DMEM probed for YB1, SYNCRIP, Flotillin-1, and CD63. <u>Middle panel:</u> Relative intensity of the YB1, SYNCRIP, Flotillin-1, and CD63 bands in the cushion density gradient fractions from the different media conditions, normalized to the band intensity of the highest SEV lane (either Fraction 6 or 7) (n=4 and 5, respectively). <u>Bottom panel:</u> Percentage of YB1, SYNCRIP, Flotillin-1 (abbreviated Flot-1), and CD63 bands found associated with Fractions 5-7, 8-10, and 11-12. Data plotted as Mean and S.E.M. *p<0.05, **p<0.01, ***p<0.001. (b) Western blot analyses of DLD-1 TCL, LEV, and DG fractions from cushion density gradients for the different media conditions probed for YB1, SYNCRIP, Flotillin-1, and CD63. <u>Middle panel:</u> Relative intensity of the YB1, SYNCRIP, Flotillin-1, and CD63 bands in the cushion density gradient fractions from the different media conditions, normalized to the band intensity of the highest SEV lane (either Fraction 6 or 7) (n=3 and 4, respectively). <u>Bottom panel:</u> Percentage of YB1, SYNCRIP, Flotillin-1 (abbreviated Flot-1), and CD63 bands found associated with Fractions 5-7, 8-10, and 11-12. Data plotted as Mean and S.E.M. *p<0.05, **p<0.01, ***p<0.001.

**Figure S6.**
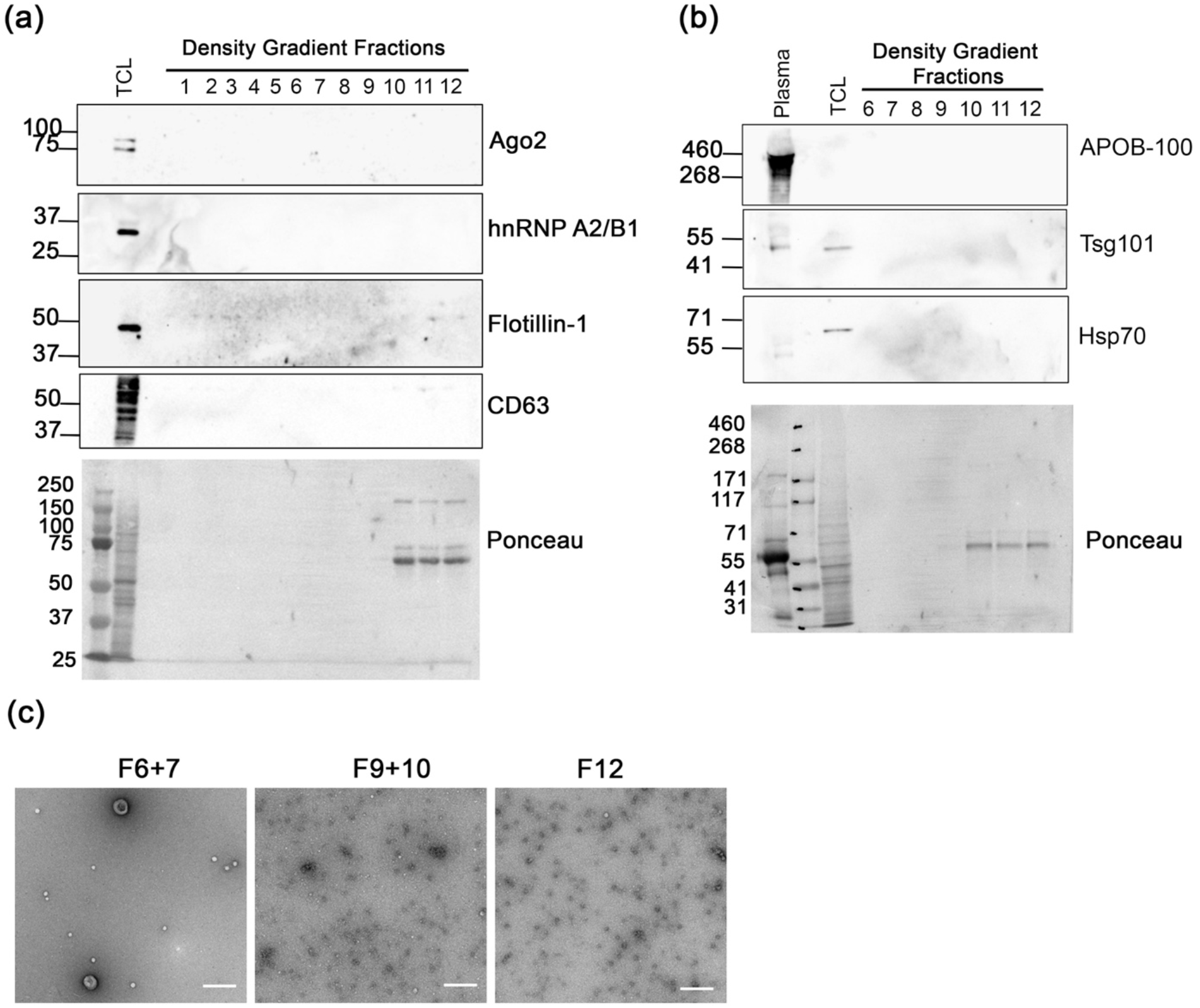
Assessment of unconditioned EV-depleted FBS in DMEM media derived from the hollow fiber bioreactor. (a) Western blot analyses of DKs-8 TCL and DG fractions from cushion density gradients for unconditioned EV-depleted FBS in DMEM derived from the hollow fiber bioreactor, probed for Ago2, hnRNP A2/B1, Flotillin-1, and CD63. (b) Western blot analyses of diluted human plasma, DKs-8 TCL, and DG fractions 6-12 for unconditioned EV-depleted FBS in DMEM derived from the hollow fiber bioreactor, probed for APOB-100, Tsg101, and Hsp70. (c) Representative TEM images of select DG fractions of unconditioned EV-depleted FBS in DMEM derived from the hollow fiber bioreactor. Scale bar shows 200 nm.

## Notes

### Competing Interest Statement

The authors have declared no competing interest.

## References

Albanese, M., Chen, Y. A., Huls, C., Gartner, K., Tagawa, T., Mejias-Perez, E., Keppler, O. T., Gobel, C., Zeidler, R., Shein, M., Schutz, A. K., & Hammerschmidt, W. (2021). MicroRNAs are minor constituents of extracellular vesicles that are rarely delivered to target cells. PLoS Genet, 17(12), e1009951. https://doi.org/10.1371/journal.pgen.1009951

Arroyo, J. D., Chevillet, J. R., Kroh, E. M., Ruf, I. K., Pritchard, C. C., Gibson, D. F., Mitchell, P. S., Bennett, C. F., Pogosova-Agadjanyan, E. L., Stirewalt, D. L., Tait, J. F., & Tewari, M. (2011). Argonaute2 complexes carry a population of circulating microRNAs independent of vesicles in human plasma. Proc Natl Acad Sci U S A, 108(12), 5003–5008. https://doi.org/10.1073/pnas.1019055108

Atwood, B. L., Woolnough, J. L., Lefevre, G. M., Ribeiro, M. S., Felsenfeld, G., & Giles, K. E. (2016). Human Argonaute 2 is tethered to ribosomal RNA through microRNA interactions. J Biol Chem. https://doi.org/10.1074/jbc.M116.725051

Auber, M., Frohlich, D., Drechsel, O., Karaulanov, E., & Kramer-Albers, E. M. (2019). Serum-free media supplements carry miRNAs that co-purify with extracellular vesicles. J Extracell Vesicles, 8(1), 1656042. https://doi.org/10.1080/20013078.2019.1656042

Aucher, A., Rudnicka, D., & Davis, D. M. (2013). MicroRNAs transfer from human macrophages to hepato-carcinoma cells and inhibit proliferation. J Immunol, 191(12), 6250–6260. https://doi.org/10.4049/jimmunol.1301728

Barman, B., Sung, B. H., Krystofiak, E., Ping, J., Ramirez, M., Millis, B., Allen, R., Prasad, N., Chetyrkin, S., Calcutt, M. W., Vickers, K., Patton, J. G., Liu, Q., & Weaver, A. M. (2022). VAP-A and its binding partner CERT drive biogenesis of RNA-containing extracellular vesicles at ER membrane contact sites. Dev Cell, 57(8), 974–994 e978. https://doi.org/10.1016/j.devcel.2022.03.012

Brennan, K., Martin, K., FitzGerald, S. P., O’Sullivan, J., Wu, Y., Blanco, A., Richardson, C., & Mc Gee, M. M. (2020). A comparison of methods for the isolation and separation of extracellular vesicles from protein and lipid particles in human serum. Sci Rep, 10(1), 1039. https://doi.org/10.1038/s41598-020-57497-7

Burroughs, A. M., Ando, Y., de Hoon, M. J., Tomaru, Y., Suzuki, H., Hayashizaki, Y., & Daub, C. O. (2011). Deep-sequencing of human Argonaute-associated small RNAs provides insight into miRNA sorting and reveals Argonaute association with RNA fragments of diverse origin. RNA Biol, 8(1), 158–177. https://doi.org/10.4161/rna.8.1.14300

Casajuana Ester, M., & Day, R. M. (2023). Production and Utility of Extracellular Vesicles with 3D Culture Methods. Pharmaceutics, 15(2). https://doi.org/10.3390/pharmaceutics15020663

Cecchin, R., Troyer, Z., Witwer, K., & Morris, K. V. (2023). Extracellular vesicles: The next generation in gene therapy delivery. Mol Ther. https://doi.org/10.1016/j.ymthe.2023.01.021

Cha, D. J., Franklin, J. L., Dou, Y., Liu, Q., Higginbotham, J. N., Demory Beckler, M., Weaver, A. M., Vickers, K., Prasad, N., Levy, S., Zhang, B., Coffey, R. J., & Patton, J. G. (2015). KRAS-dependent sorting of miRNA to exosomes. Elife, 4, e07197. https://doi.org/10.7554/eLife.07197

Chen, W. X., Liu, X. M., Lv, M. M., Chen, L., Zhao, J. H., Zhong, S. L., Ji, M. H., Hu, Q., Luo, Z., Wu, J. Z., & Tang, J. H. (2014). Exosomes from drug-resistant breast cancer cells transmit chemoresistance by a horizontal transfer of microRNAs. PLoS One, 9(4), e95240. https://doi.org/10.1371/journal.pone.0095240

Chiou, N. T., Kageyama, R., & Ansel, K. M. (2018). Selective Export into Extracellular Vesicles and Function of tRNA Fragments during T Cell Activation. Cell Rep, 25(12), 3356–3370 e3354. https://doi.org/10.1016/j.celrep.2018.11.073

Demory Beckler, M., Higginbotham, J. N., Franklin, J. L., Ham, A. J., Halvey, P. J., Imasuen, I. E., Whitwell, C., Li, M., Liebler, D. C., & Coffey, R. J. (2013). Proteomic analysis of exosomes from mutant KRAS colon cancer cells identifies intercellular transfer of mutant KRAS. Mol Cell Proteomics, 12(2), 343–355. https://doi.org/10.1074/mcp.M112.022806

Dhondt, B., Rousseau, Q., De Wever, O., & Hendrix, A. (2016). Function of extracellular vesicle-associated miRNAs in metastasis. Cell Tissue Res. https://doi.org/10.1007/s00441-016-2430-x

Dixson, A. C., Dawson, T. R., Di Vizio, D., & Weaver, A. M. (2023). Context-specific regulation of extracellular vesicle biogenesis and cargo selection. Nat Rev Mol Cell Biol. https://doi.org/10.1038/s41580-023-00576-0

Driedonks, T. A. P., Nijen Twilhaar, M. K., & Nolte-’t Hoen, E. N. M. (2019). Technical approaches to reduce interference of Fetal calf serum derived RNA in the analysis of extracellular vesicle RNA from cultured cells. J Extracell Vesicles, 8(1), 1552059. https://doi.org/10.1080/20013078.2018.1552059

Dupuis-Sandoval, F., Poirier, M., & Scott, M. S. (2015). The emerging landscape of small nucleolar RNAs in cell biology. Wiley Interdiscip Rev RNA, 6(4), 381–397. https://doi.org/10.1002/wrna.1284

Erickson, H. P. (2009). Size and shape of protein molecules at the nanometer level determined by sedimentation, gel filtration, and electron microscopy. Biol Proced Online, 11, 32–51. https://doi.org/10.1007/s12575-009-9008-x

Gardiner, C., Ferreira, Y. J., Dragovic, R. A., Redman, C. W., & Sargent, I. L. (2013). Extracellular vesicle sizing and enumeration by nanoparticle tracking analysis. J Extracell Vesicles, 2. https://doi.org/10.3402/jev.v2i0.19671

Geekiyanage, H., Rayatpisheh, S., Wohlschlegel, J. A., Brown, R., Jr., & Ambros, V. (2020). Extracellular microRNAs in human circulation are associated with miRISC complexes that are accessible to anti-AGO2 antibody and can bind target mimic oligonucleotides. Proc Natl Acad Sci U S A, 117(39), 24213–24223. https://doi.org/10.1073/pnas.2008323117

Han, M., Yang, H., Lu, X., Li, Y., Liu, Z., Li, F., Shang, Z., Wang, X., Li, X., Li, J., Liu, H., & Xin, T. (2022). Three-Dimensional-Cultured MSC-Derived Exosome-Hydrogel Hybrid Microneedle Array Patch for Spinal Cord Repair. Nano Lett, 22(15), 6391–6401. https://doi.org/10.1021/acs.nanolett.2c02259

Haussecker, D., Huang, Y., Lau, A., Parameswaran, P., Fire, A. Z., & Kay, M. A. (2010). Human tRNA-derived small RNAs in the global regulation of RNA silencing. Rna, 16(4), 673–695. https://doi.org/10.1261/rna.2000810

Hobor, F., Dallmann, A., Ball, N. J., Cicchini, C., Battistelli, C., Ogrodowicz, R. W., Christodoulou, E., Martin, S. R., Castello, A., Tripodi, M., Taylor, I. A., & Ramos, A. (2018). A cryptic RNA-binding domain mediates Syncrip recognition and exosomal partitioning of miRNA targets. Nat Commun, 9(1), 831. https://doi.org/10.1038/s41467-018-03182-3

Jeppesen, D. K., Fenix, A. M., Franklin, J. L., Higginbotham, J. N., Zhang, Q., Zimmerman, L. J., Liebler, D. C., Ping, J., Liu, Q., Evans, R., Fissell, W. H., Patton, J. G., Rome, L. H., Burnette, D. T., & Coffey, R. J. (2019). Reassessment of Exosome Composition. Cell, 177(2), 428–445 e418. https://doi.org/10.1016/j.cell.2019.02.029

Jiang, Y., Qiu, Q., Jing, X., Song, Z., Zhang, Y., Wang, C., Liu, K., Ye, F., Ji, X., Luo, F., & Zhao, R. (2023). Cancer-associated fibroblast-derived exosome miR-181b-3p promotes the occurrence and development of colorectal cancer by regulating SNX2 expression. Biochem Biophys Res Commun, 641, 177–185. https://doi.org/10.1016/j.bbrc.2022.12.026

Kishore, S., Gruber, A. R., Jedlinski, D. J., Syed, A. P., Jorjani, H., & Zavolan, M. (2013). Insights into snoRNA biogenesis and processing from PAR-CLIP of snoRNA core proteins and small RNA sequencing. Genome Biol, 14(5), R45. https://doi.org/10.1186/gb-2013-14-5-r45

Kornilov, R., Puhka, M., Mannerstrom, B., Hiidenmaa, H., Peltoniemi, H., Siljander, P., Seppanen-Kaijansinkko, R., & Kaur, S. (2018). Efficient ultrafiltration-based protocol to deplete extracellular vesicles from fetal bovine serum. J Extracell Vesicles, 7(1), 1422674. https://doi.org/10.1080/20013078.2017.1422674

Kossinova, O. A., Gopanenko, A. V., Tamkovich, S. N., Krasheninina, O. A., Tupikin, A. E., Kiseleva, E., Yanshina, D. D., Malygin, A. A., Ven’yaminova, A. G., Kabilov, M. R., & Karpova, G. G. (2017). Cytosolic YB-1 and NSUN2 are the only proteins recognizing specific motifs present in mRNAs enriched in exosomes. Biochim Biophys Acta Proteins Proteom, 1865(6), 664–673. https://doi.org/10.1016/j.bbapap.2017.03.010

Lai, C. P., Kim, E. Y., Badr, C. E., Weissleder, R., Mempel, T. R., Tannous, B. A., & Breakefield, X. O. (2015). Visualization and tracking of tumour extracellular vesicle delivery and RNA translation using multiplexed reporters. Nat Commun, 6, 7029. https://doi.org/10.1038/ncomms8029

Larios, J., Mercier, V., Roux, A., & Gruenberg, J. (2020). ALIX- and ESCRT-III-dependent sorting of tetraspanins to exosomes. J Cell Biol, 219(3). https://doi.org/10.1083/jcb.201904113

Lehrich, B. M., Liang, Y., Khosravi, P., Federoff, H. J., & Fiandaca, M. S. (2018). Fetal Bovine Serum-Derived Extracellular Vesicles Persist within Vesicle-Depleted Culture Media. Int J Mol Sci, 19(11). https://doi.org/10.3390/ijms19113538

Leidal, A. M., Huang, H. H., Marsh, T., Solvik, T., Zhang, D., Ye, J., Kai, F., Goldsmith, J., Liu, J. Y., Huang, Y. H., Monkkonen, T., Vlahakis, A., Huang, E. J., Goodarzi, H., Yu, L., Wiita, A. P., & Debnath, J. (2020). The LC3-conjugation machinery specifies the loading of RNA-binding proteins into extracellular vesicles. Nat Cell Biol, 22(2), 187–199. https://doi.org/10.1038/s41556-019-0450-y

Li, C., Ni, Y. Q., Xu, H., Xiang, Q. Y., Zhao, Y., Zhan, J. K., He, J. Y., Li, S., & Liu, Y. S. (2021). Roles and mechanisms of exosomal non-coding RNAs in human health and diseases. Signal Transduct Target Ther, 6(1), 383. https://doi.org/10.1038/s41392-021-00779-x

Li, K., Wong, D. K., Hong, K. Y., & Raffai, R. L. (2018). Cushioned-Density Gradient Ultracentrifugation (C-DGUC): A Refined and High Performance Method for the Isolation, Characterization, and Use of Exosomes. Methods Mol Biol, 1740, 69–83. https://doi.org/10.1007/978-1-4939-7652-2_7

Liao, J., Liu, R., Shi, Y. J., Yin, L. H., & Pu, Y. P. (2016). Exosome-shuttling microRNA-21 promotes cell migration and invasion-targeting PDCD4 in esophageal cancer. Int J Oncol, 48(6), 2567–2579. https://doi.org/10.3892/ijo.2016.3453

Liu, X.-M., Ma, L., & Schekman, R. (2021). Selective sorting of microRNAs into exosomes by phase-separated YBX1 condensates. eLife, 10. https://doi.org/10.7554/eLife.71982

Lunavat, T. R., Cheng, L., Kim, D. K., Bhadury, J., Jang, S. C., Lässer, C., Sharples, R. A., López, M. D., Nilsson, J., Gho, Y. S., Hill, A. F., & Lötvall, J. (2015). Small RNA deep sequencing discriminates subsets of extracellular vesicles released by melanoma cells--Evidence of unique microRNA cargos. RNA Biol, 12(8), 810–823. https://doi.org/10.1080/15476286.2015.1056975

Maas, S. L. N., Breakefield, X. O., & Weaver, A. M. (2017). Extracellular Vesicles: Unique Intercellular Delivery Vehicles. Trends Cell Biol, 27(3), 172–188. https://doi.org/10.1016/j.tcb.2016.11.003

Mannerstrom, B., Paananen, R. O., Abu-Shahba, A. G., Moilanen, J., Seppanen-Kaijansinkko, R., & Kaur, S. (2019). Extracellular small non-coding RNA contaminants in fetal bovine serum and serum-free media. Sci Rep, 9(1), 5538. https://doi.org/10.1038/s41598-019-41772-3

Mantel, P. Y., Hjelmqvist, D., Walch, M., Kharoubi-Hess, S., Nilsson, S., Ravel, D., Ribeiro, M., Gruring, C., Ma, S., Padmanabhan, P., Trachtenberg, A., Ankarklev, J., Brancucci, N. M., Huttenhower, C., Duraisingh, M. T., Ghiran, I., Kuo, W. P., Filgueira, L., Martinelli, R., & Marti, M. (2016). Infected erythrocyte-derived extracellular vesicles alter vascular function via regulatory Ago2-miRNA complexes in malaria. Nat Commun, 7, 12727. https://doi.org/10.1038/ncomms12727

McKenzie, A. J., Hoshino, D., Hong, N. H., Cha, D. J., Franklin, J. L., Coffey, R. J., Patton, J. G., & Weaver, A. M. (2016). KRAS-MEK Signaling Controls Ago2 Sorting into Exosomes. Cell Rep, 15(5), 978–987. https://doi.org/10.1016/j.celrep.2016.03.085

Meister, G. (2013). Argonaute proteins: functional insights and emerging roles. Nat Rev Genet, 14(7), 447–459. https://doi.org/10.1038/nrg3462

Murphy, D. E., de Jong, O. G., Brouwer, M., Wood, M. J., Lavieu, G., Schiffelers, R. M., & Vader, P. (2019). Extracellular vesicle-based therapeutics: natural versus engineered targeting and trafficking. Exp Mol Med, 51(3), 1–12. https://doi.org/10.1038/s12276-019-0223-5

O’Brien, K., Breyne, K., Ughetto, S., Laurent, L. C., & Breakefield, X. O. (2020). RNA delivery by extracellular vesicles in mammalian cells and its applications. Nat Rev Mol Cell Biol, 21(10), 585–606. https://doi.org/10.1038/s41580-020-0251-y

Oszvald, A., Szvicsek, Z., Papai, M., Kelemen, A., Varga, Z., Tolgyes, T., Dede, K., Bursics, A., Buzas, E. I., & Wiener, Z. (2020). Fibroblast-Derived Extracellular Vesicles Induce Colorectal Cancer Progression by Transmitting Amphiregulin. Front Cell Dev Biol, 8, 558. https://doi.org/10.3389/fcell.2020.00558

Patel, M. R., & Weaver, A. M. (2021). Astrocyte-derived small extracellular vesicles promote synapse formation via fibulin-2-mediated TGF-beta signaling. Cell Rep, 34(10), 108829. https://doi.org/10.1016/j.celrep.2021.108829

Patton, J. G., Franklin, J. L., Weaver, A. M., Vickers, K., Zhang, B., Coffey, R. J., Ansel, K. M., Blelloch, R., Goga, A., Huang, B., L’Etoille, N., Raffai, R. L., Lai, C. P., Krichevsky, A. M., Mateescu, B., Greiner, V. J., Hunter, C., Voinnet, O., & McManus, M. T. (2015). Biogenesis, delivery, and function of extracellular RNA. J Extracell Vesicles, 4, 27494. https://doi.org/10.3402/jev.v4.27494

Persson, H., Kvist, A., Vallon-Christersson, J., Medstrand, P., Borg, A., & Rovira, C. (2009). The non-coding RNA of the multidrug resistance-linked vault particle encodes multiple regulatory small RNAs. Nat Cell Biol, 11(10), 1268–1271. https://doi.org/10.1038/ncb1972

Pham, C. V., Midge, S., Barua, H., Zhang, Y., Ngoc-Gia Nguyen, T., Barrero, R. A., Duan, A., Yin, W., Jiang, G., Hou, Y., Zhou, S., Wang, Y., Xie, X., Tran, P. H. L., Xiang, D., & Duan, W. (2021). Bovine extracellular vesicles contaminate human extracellular vesicles produced in cell culture conditioned medium when ‘exosome-depleted serum’ is utilised. Arch Biochem Biophys, 708, 108963. https://doi.org/10.1016/j.abb.2021.108963

Sahoo, S., Klychko, E., Thorne, T., Misener, S., Schultz, K. M., Millay, M., Ito, A., Liu, T., Kamide, C., Agrawal, H., Perlman, H., Qin, G., Kishore, R., & Losordo, D. W. (2011). Exosomes from human CD34(+) stem cells mediate their proangiogenic paracrine activity. Circ Res, 109(7), 724–728. https://doi.org/10.1161/circresaha.111.253286

Santangelo, L., Giurato, G., Cicchini, C., Montaldo, C., Mancone, C., Tarallo, R., Battistelli, C., Alonzi, T., Weisz, A., & Tripodi, M. (2016). The RNA-Binding Protein SYNCRIP Is a Component of the Hepatocyte Exosomal Machinery Controlling MicroRNA Sorting. Cell Rep, 17(3), 799–808. https://doi.org/10.1016/j.celrep.2016.09.031

Sato, S., Vasaikar, S., Eskaros, A., Kim, Y., Lewis, J. S., Zhang, B., Zijlstra, A., & Weaver, A. M. (2019). EPHB2 carried on small extracellular vesicles induces tumor angiogenesis via activation of ephrin reverse signaling. JCI Insight, 4(23). https://doi.org/10.1172/jci.insight.132447

Sato, S., & Weaver, A. M. (2018). Extracellular vesicles: important collaborators in cancer progression. Essays Biochem, 62(2), 149–163. https://doi.org/10.1042/EBC20170080

Schmittgen, T. D., & Livak, K. J. (2008). Analyzing real-time PCR data by the comparative C(T) method. Nat Protoc, 3(6), 1101–1108. https://doi.org/10.1038/nprot.2008.73

Schneider, C. A., Rasband, W. S., & Eliceiri, K. W. (2012). NIH Image to ImageJ: 25 years of image analysis. Nat Methods, 9(7), 671–675. https://doi.org/10.1038/nmeth.2089

Shelke, G. V., Lasser, C., Gho, Y. S., & Lotvall, J. (2014). Importance of exosome depletion protocols to eliminate functional and RNA-containing extracellular vesicles from fetal bovine serum. J Extracell Vesicles, 3. https://doi.org/10.3402/jev.v3.24783

Shirasawa, S., Furuse, M., Yokoyama, N., & Sasazuki, T. (1993). Altered Growth of Human Colon Cancer Cell-Lines Disrupted at Activated Ki-Ras. Science, 260(5104), 85–88. https://doi.org/10.1126/science.8465203

Shurtleff, M., Karfilis, K. V., Temoche-Diaz, M., Ri, S., & Schekman, R. (2016). Y-box protein 1 is required to sort microRNAs into exosomes in cells and in a cell-free reaction. https://doi.org/10.1101/040238

Sung, B. H., Ketova, T., Hoshino, D., Zijlstra, A., & Weaver, A. M. (2015). Directional cell movement through tissues is controlled by exosome secretion. Nat Commun, 6, 7164. https://doi.org/10.1038/ncomms8164

Thery, C., Amigorena, S., Raposo, G., & Clayton, A. (2006). Isolation and characterization of exosomes from cell culture supernatants and biological fluids. Curr Protoc Cell Biol, Chapter 3, Unit 3 22. https://doi.org/10.1002/0471143030.cb0322s30

Thippabhotla, S., Zhong, C., & He, M. (2019). 3D cell culture stimulates the secretion of in vivo like extracellular vesicles. Sci Rep, 9(1), 13012. https://doi.org/10.1038/s41598-019-49671-3

Tosar, J. P., Cayota, A., Eitan, E., Halushka, M. K., & Witwer, K. W. (2017). Ribonucleic artefacts: are some extracellular RNA discoveries driven by cell culture medium components? J Extracell Vesicles, 6(1), 1272832. https://doi.org/10.1080/20013078.2016.1272832

Tu, J., Luo, X., Liu, H., Zhang, J., & He, M. (2021). Cancer spheroids derived exosomes reveal more molecular features relevant to progressed cancer. Biochem Biophys Rep, 26, 101026. https://doi.org/10.1016/j.bbrep.2021.101026

Turchinovich, A., Weiz, L., Langheinz, A., & Burwinkel, B. (2011). Characterization of extracellular circulating microRNA. Nucleic Acids Res, 39(16), 7223–7233. https://doi.org/10.1093/nar/gkr254

Urzi, O., Bagge, R. O., & Crescitelli, R. (2022). The dark side of foetal bovine serum in extracellular vesicle studies. J Extracell Vesicles, 11(10), e12271. https://doi.org/10.1002/jev2.12271

van Balkom, B. W., Eisele, A. S., Pegtel, D. M., Bervoets, S., & Verhaar, M. C. (2015). Quantitative and qualitative analysis of small RNAs in human endothelial cells and exosomes provides insights into localized RNA processing, degradation and sorting. J Extracell Vesicles, 4, 26760. https://doi.org/10.3402/jev.v4.26760

Vickers, K. C., Palmisano, B. T., Shoucri, B. M., Shamburek, R. D., & Remaley, A. T. (2011). MicroRNAs are transported in plasma and delivered to recipient cells by high-density lipoproteins. Nat Cell Biol, 13(4), 423–433. https://doi.org/10.1038/ncb2210

Villarroya-Beltri, C., Gutierrez-Vazquez, C., Sanchez-Cabo, F., Perez-Hernandez, D., Vazquez, J., Martin-Cofreces, N., Martinez-Herrera, D. J., Pascual-Montano, A., Mittelbrunn, M., & Sanchez-Madrid, F. (2013). Sumoylated hnRNPA2B1 controls the sorting of miRNAs into exosomes through binding to specific motifs. Nat Commun, 4, 2980. https://doi.org/10.1038/ncomms3980

Weaver, A. M., & Patton, J. G. (2020). Argonautes in Extracellular Vesicles: Artifact or Selected Cargo? Cancer Res, 80(3), 379–381. https://doi.org/10.1158/0008-5472.CAN-19-2782

Wei, Z., Batagov, A. O., Carter, D. R., & Krichevsky, A. M. (2016). Fetal Bovine Serum RNA Interferes with the Cell Culture derived Extracellular RNA. Sci Rep, 6, 31175. https://doi.org/10.1038/srep31175

Woolnough, J. L., Atwood, B. L., & Giles, K. E. (2015). Argonaute 2 Binds Directly to tRNA Genes and Promotes Gene Repression in cis. Mol Cell Biol, 35(13), 2278–2294. https://doi.org/10.1128/mcb.00076-15

Yuan, X., Sun, L., Jeske, R., Nkosi, D., York, S. B., Liu, Y., Grant, S. C., Meckes, D. G., Jr., & Li, Y. (2022). Engineering extracellular vesicles by three-dimensional dynamic culture of human mesenchymal stem cells. J Extracell Vesicles, 11(6), e12235. https://doi.org/10.1002/jev2.12235

Zand Karimi, H., Baldrich, P., Rutter, B. D., Borniego, L., Zajt, K. K., Meyers, B. C., & Innes, R. W. (2022). Arabidopsis apoplastic fluid contains sRNA- and circular RNA-protein complexes that are located outside extracellular vesicles. Plant Cell, 34(5), 1863–1881. https://doi.org/10.1093/plcell/koac043

Zhang, Q., Jeppesen, D. K., Higginbotham, J. N., Franklin, J. L., & Coffey, R. J. (2023). Comprehensive isolation of extracellular vesicles and nanoparticles. Nat Protoc, 18(5), 1462–1487. https://doi.org/10.1038/s41596-023-00811-0

Zhang, Q., Jeppesen, D. K., Higginbotham, J. N., Graves-Deal, R., Trinh, V. Q., Ramirez, M. A., Sohn, Y., Neininger, A. C., Taneja, N., McKinley, E. T., Niitsu, H., Cao, Z., Evans, R., Glass, S. E., Ray, K. C., Fissell, W. H., Hill, S., Rose, K. L., Huh, W. J., . . . Coffey, R. J. (2021). Supermeres are functional extracellular nanoparticles replete with disease biomarkers and therapeutic targets. Nat Cell Biol, 23(12), 1240–1254. https://doi.org/10.1038/s41556-021-00805-8

Zhang, Y., Liu, D., Chen, X., Li, J., Li, L., Bian, Z., Sun, F., Lu, J., Yin, Y., Cai, X., Sun, Q., Wang, K., Ba, Y., Wang, Q., Wang, D., Yang, J., Liu, P., Xu, T., Yan, Q., . . . Zhang, C. Y. (2010). Secreted monocytic miR-150 enhances targeted endothelial cell migration. Mol Cell, 39(1), 133–144. https://doi.org/10.1016/j.molcel.2010.06.010

Zheng, X., Baker, H., Hancock, W. S., Fawaz, F., McCaman, M., & Pungor, E., Jr. (2006). Proteomic analysis for the assessment of different lots of fetal bovine serum as a raw material for cell culture. Part IV. Application of proteomics to the manufacture of biological drugs. Biotechnol Prog, 22(5), 1294–1300. https://doi.org/10.1021/bp060121o

Zhou, X. S., Wang, L., Zou, W. X., Chen, X. Q., Roizman, B., & Zhou, G. G. (2020). hnRNPA2B1 Associated with Recruitment of RNA into Exosomes Plays a Key Role in Herpes Simplex Virus 1 Release from Infected Cells. Journal of Virology, 94(13). https://doi.org/10.1128/JVI.00367-20

